# Long-read assemblies reveal structural diversity in genomes of organelles - an example with *Acacia pycnantha*

**DOI:** 10.1101/2020.12.22.423164

**Authors:** Anna E. Syme, Todd G.B. McLay, Frank Udovicic, David J. Cantrill, Daniel J. Murphy

**Author notes:** Corresponding Author: Anna Syme.

## Abstract

Although organelle genomes are typically represented as single, static, circular molecules, there is evidence that the chloroplast genome exists in two structural haplotypes and that the mitochondrial genome can display multiple circular, linear or branching forms. We sequenced and assembled chloroplast and mitochondrial genomes of the Golden Wattle, *Acacia pycnantha,* using long reads, iterative baiting to extract organelle-only reads, and several assembly algorithms to explore genomic structure. Using a *de novo* assembly approach agnostic to previous hypotheses about structure, we found different assemblies revealed contrasting arrangements of genomic segments; a hypothesis supported by mapped reads spanning alternate paths.

## Introduction

Genomes from organelles such as those of the chloroplast and mitochondria have predominantly been sequenced by technologies producing read lengths of between 75-300 base pairs. These are considered “short” reads by comparison with newer technologies that can routinely produce “long” reads of 10,000 base pairs and longer.

Although some of the earlier organelle genome assemblies were inferred from long range PCR and associated strategies (Dierckxsens, Mardulyn, and Smits 2017), the bulk of existing assemblies are based on short-read sequencing data which was comparatively easier and cheaper to obtain.

Short-read assemblies are challenging, as repeats longer than read length can’t be unambiguously placed within the genome assembly. This is evident in chloroplast genome assemblies, where the inverted repeats (IR) are typically assembled into a single contig, and are then manually duplicated in the assembly result to recreate the circular structure. There are many reasons that this is less than ideal, one being that variation in repeats may not be captured, and the IR boundaries are not always reconstructed accurately.

Short-read assemblies have additional challenges for mitochondrial genomes, due to their larger size and structural complexity. Whereas the chloroplast genome (plastome) in land plants is typically ~160 Kbp and circular, the plant mitochondrial genome (mitome) is ~800 Kbp and likely exists in multiple dynamic structures. Although the mitome has traditionally been represented as a single circular structure, there is physical evidence of multiple shapes (Bendich 1996). Long sequencing reads indicate several linear, branched, or smaller circular structures (Kozik et al. 2019; Jackman et al. 2020) that may recombine at repeat regions (Z. Chen et al. 2017).

Long sequencing reads may be able to span repetitive regions (depending on length) and should better capture their placement in assemblies, thereby revealing more of the structural complexity in both plastomes and mitomes. Presently, long reads from technologies such as Oxford Nanopore and PacBio have a higher error-rate than short reads from Illumina. Thus, reads from both can be combined when assembling genomes: long reads can reveal broad organelle genome structure, and short reads can correct errors. This hybrid approach has demonstrated improved accuracy in organelle genome assembly (Wang et al. 2018).

Recent work using these combined technologies has been highly successful in revealing new things about organelle genome structure. Long reads provide strong evidence that the plastome exists in two structural haplotypes in equal proportions across land plants (Wang and Lanfear 2019) which supports certain theories of recombination. Gymnosperm mitome assemblies based on long reads reveal significant complexity in mitome structure, and branching may be related to recombination processes (Jackman et al. 2020).

In this paper we used long (Oxford Nanopore) and short (Illumina) sequencing reads to assemble the plastome and mitome of *Acacia pycnantha* (Golden Wattle, Australia’s floral emblem). The genus *Acacia* has more than 1000 species and is iconic and economically important, yet currently lacks any long-read organelle assemblies. Data from nuclear ribosomal DNA and plastomes have provided a good basis for phylogenetic investigation (Williams et al. 2016). Three plastomes and one mitome for the genus are available in RefSeq (O’Leary et al. 2016), and 94 additional partial assemblies (incomplete with gaps in non-coding, repeat rich regions) in NCBI, many from Williams et al. (2006). These new long-read assemblies will complement and expand our knowledge of *Acacia* organelle structures.

To further facilitate exploration of genome structure and to limit introducing bias or errors, we assembled sequencing reads *de novo*, rather than mapping them to an existing assembly. Analysis steps are automated in reproducible scripts with freely available tools.

## Methods

### Analysis rationale

#### Obtaining organelle reads

One of the most difficult parts of this analysis was to extract the organelle-only reads from the full genomic read sets that contain a mix of nuclear, mitome and plastome reads. We used an iterative approach to do this. First, gene coding sequences from related *Acacia* species were used as baits to extract organelle Nanopore reads. These reads were assembled, and the assembly itself used as bait for repeat organelle read extraction from the full Nanopore read set. These reads were assembled, and this second assembly was used as bait to extract the short Illumina reads from the full Illumina read set. The short reads were then used to polish the assembly (Figure 1). Additional assemblies were completed in different configurations as discussed in more detail below.

**Figure 1.**
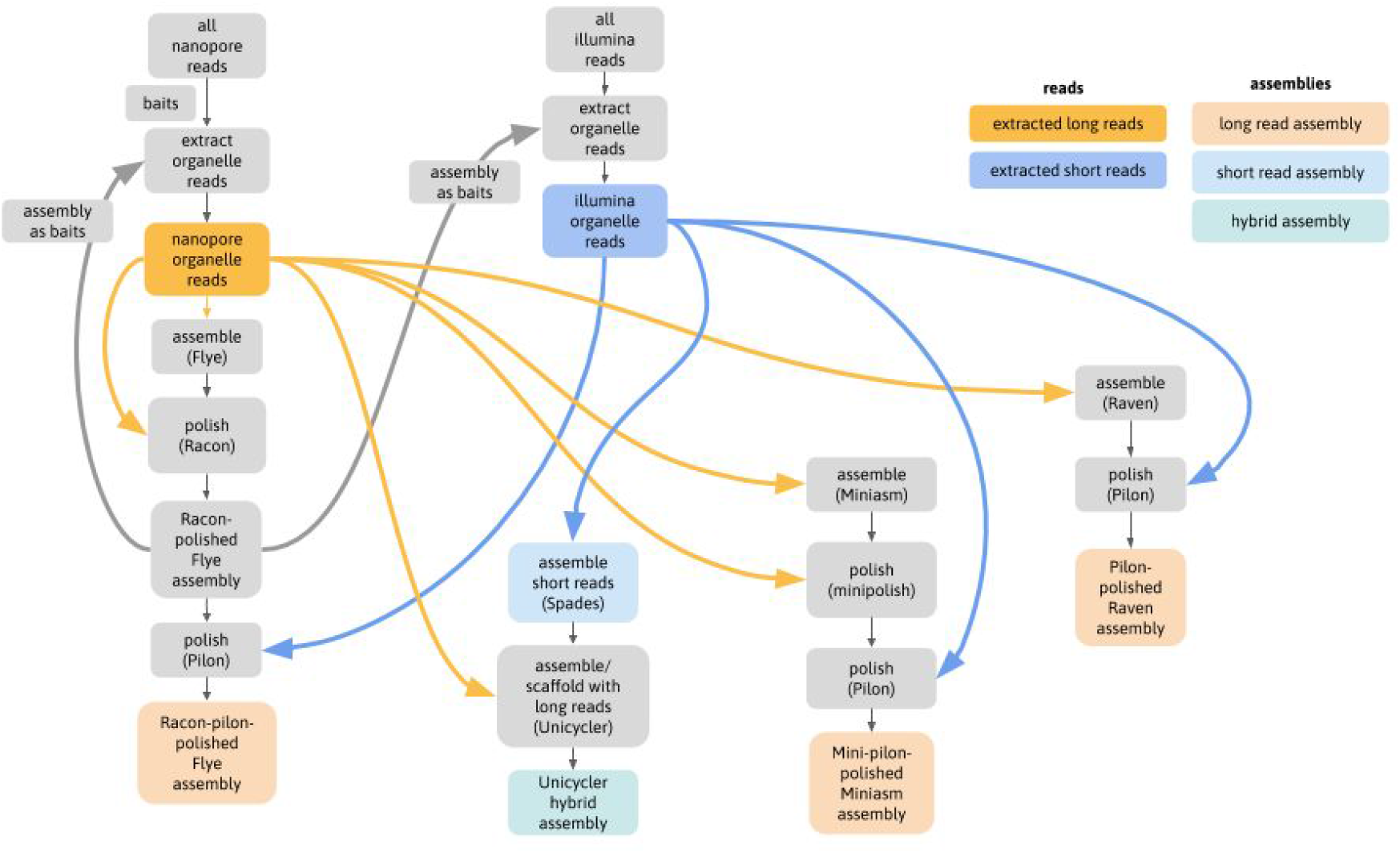
Analysis flowchart. Main steps in the analysis, showing use of initial assembly as baits for further read extraction, and use of read sets in assembly polishing steps.

#### Assembling organelle genomes

As short-read technologies have provided the dominant source of organelle sequencing read data for the past ten years, assembly tools have by necessity also been based on reads of these lengths and fidelity, and typically relied on mapping the reads to existing assemblies. Over the past four years, the tools NOVOPlasty (Dierckxsens, Mardulyn, and Smits 2017) and GetOrganelle (Jin et al. 2019) have been developed using iterative baiting algorithms to perform *de novo* organelle assemblies, demonstrating good success at improving accuracy. However, they are not configured to work with long read data, which differs not only in read length, but read length variability and a higher error profile (Rang, Kloosterman, and de Ridder 2018). In this analysis, where we specifically wanted to investigate assemblies based on long reads, we therefore used assemblers that have been optimised for this type of data.

#### Choice of assembly tools

Due to a likely shared ancestry, organelle genomes have similarity to bacterial genomes (McFadden 1999). It is therefore appropriate to consider methods used for bacterial assembly, an area of active research because accurate assemblies underpin many areas of public health. A recent benchmarking of bacterial assembly tools for long reads found the best performers to be Flye (Kolmogorov et al. 2019), Raven (Vaser and Šikić 2020) and Miniasm (Li 2016) + Minipolish (R. R. Wick and Holt 2019), but no single tool performed best on all metrics such as reliability, circularisation, errors, and completeness (R. R. Wick and Holt 2019).

Here, we chose to use a combination of these well-tested assemblers, which also capture the diversity of algorithms in current use. For example, Flye uses approximate repeat graphs; Miniasm is a true Overlap-Layout-Consensus (OLC) method and only outputs unitigs; and Raven combines an OLC method with improved graph cleaning by removing unsupported overlaps. All are designed to work with ‘noisy’ long reads such as Oxford Nanopore sequences. Because we are also including short reads in this analysis, we added an assembly by the tool Unicycler (R. R. Wick et al. 2017) - a hybrid method using an initial short-read assembly followed by long-read scaffolding (Figure 1).

#### Annotations

This analysis is primarily concerned with establishing first-pass assemblies for *Acacia* organelles, using long reads to explore structural configuration. Annotation is the next step to further investigate gene and feature content, arrangement, loss or duplication, transfer between organelles and nuclear genomes, and comparison with related and more distant species. In appreciation of the complexity of this and the need for domain-specific knowledge to best produce a useful annotation, this work does not attempt to present a completed or final annotation of these organelle genomes. However, basic annotations are presented to provide an initial visualisation of the feature landscape of these organelles. We used the tool GeSeq (Tillich et al. 2017) to produce annotations, which is automated, reproducible, and has been used successfully for plant mitomes and plastomes (Frommer et al. 2020; Guyeux et al. 2019). Future editing and refinements of these annotations are therefore expected and will no doubt provide improvement in interpretation of structural and physiological processes. In fact, a continuous iterative process of refinement of assemblies, then annotations, then assemblies, and so on, would be beneficial, particularly in the study of non-model organisms.

### Sample and sequencing

Young leaves were collected from a Golden Wattle, *Acacia pycnantha*, in the Australian National Botanic Gardens (voucher details: CANB 748486 S.R. Donaldson 3550 12/10/2007) and DNA was extracted from fresh tissue (McLay n.d.). Sequencing was performed by the Australian Genome Research Facility (Melbourne, Australia). Oxford Nanopore sequencing used a PromethION R9.4.1 flow cell and basecalling with Guppy v. 3.2.4, producing ~5.5 million reads, longest ~170 Kbp, total ~60 Gbp (Table 1). Illumina sequencing used a TruSeq DNA Nano protocol, a NovaSeq 6000 S4 flow cell, basecalling with Illumina RTA v3.4.4, de-multiplexing and FASTQ file generation with Illumina bcl2fastq pipeline v. 2.20.0.422, producing ~480 million 300 bp read pairs, totalling ~140 Gbp (Table 1).

**Table 1.**
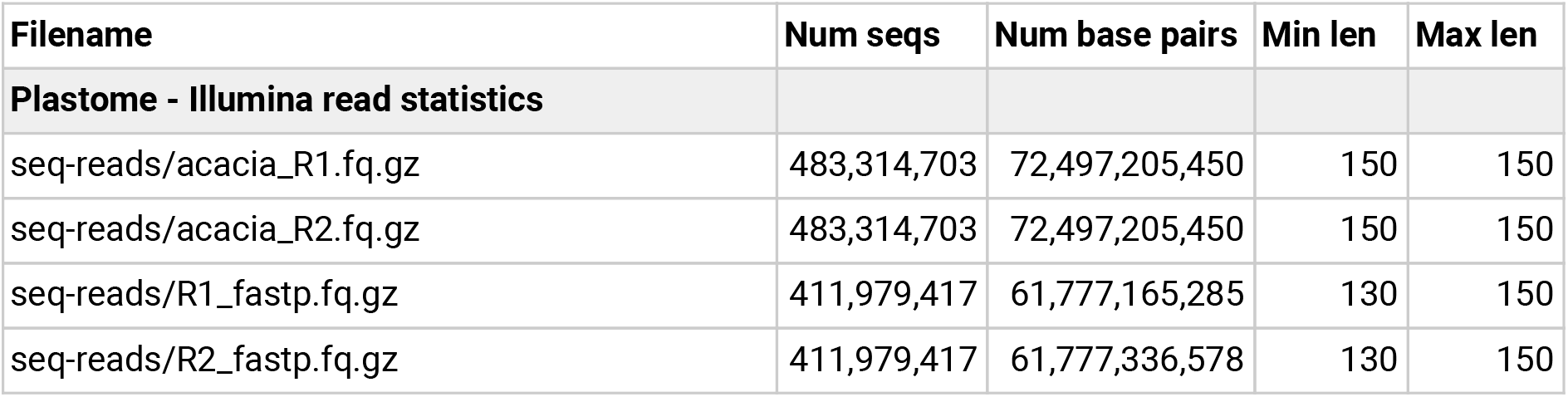
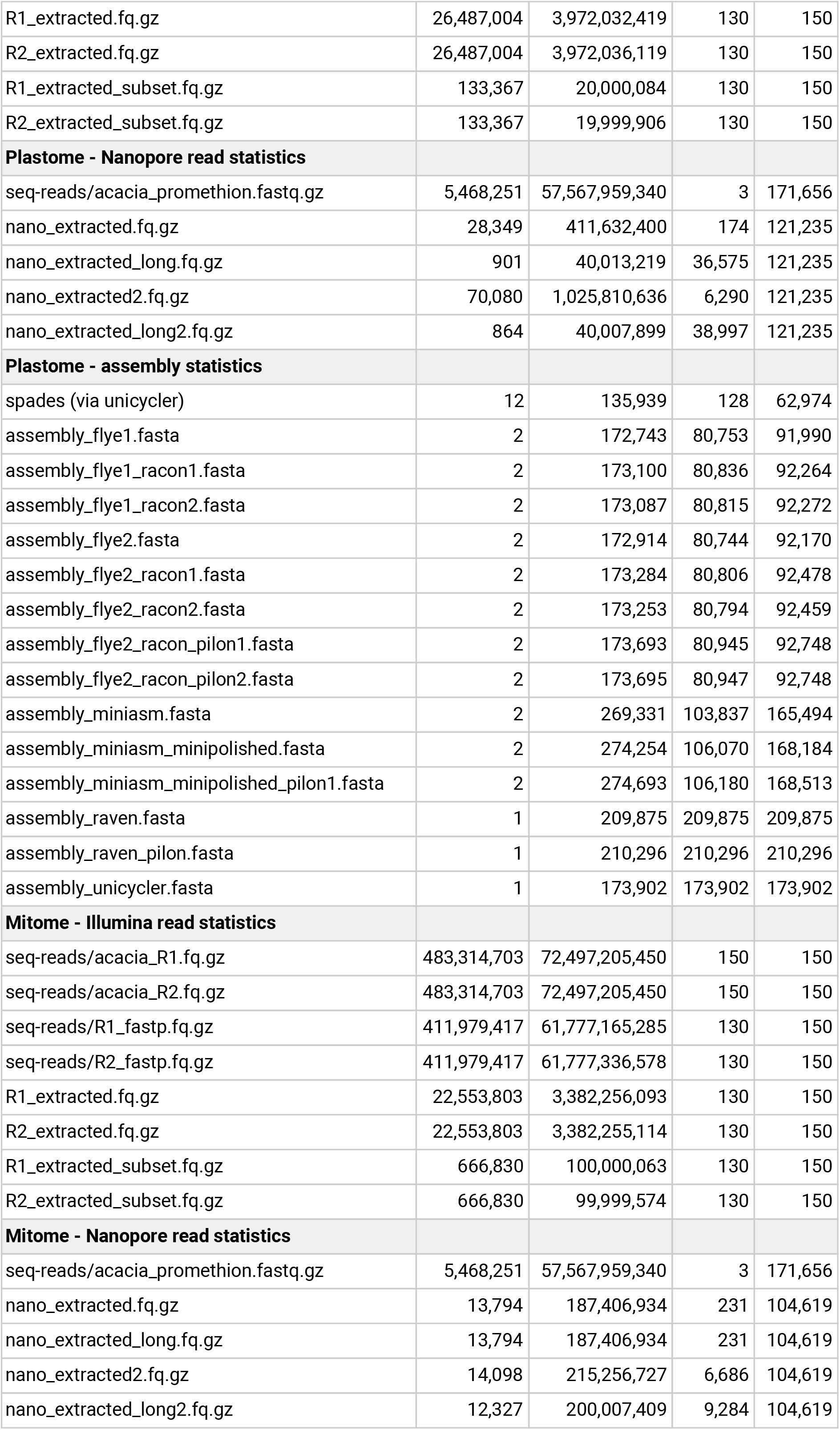
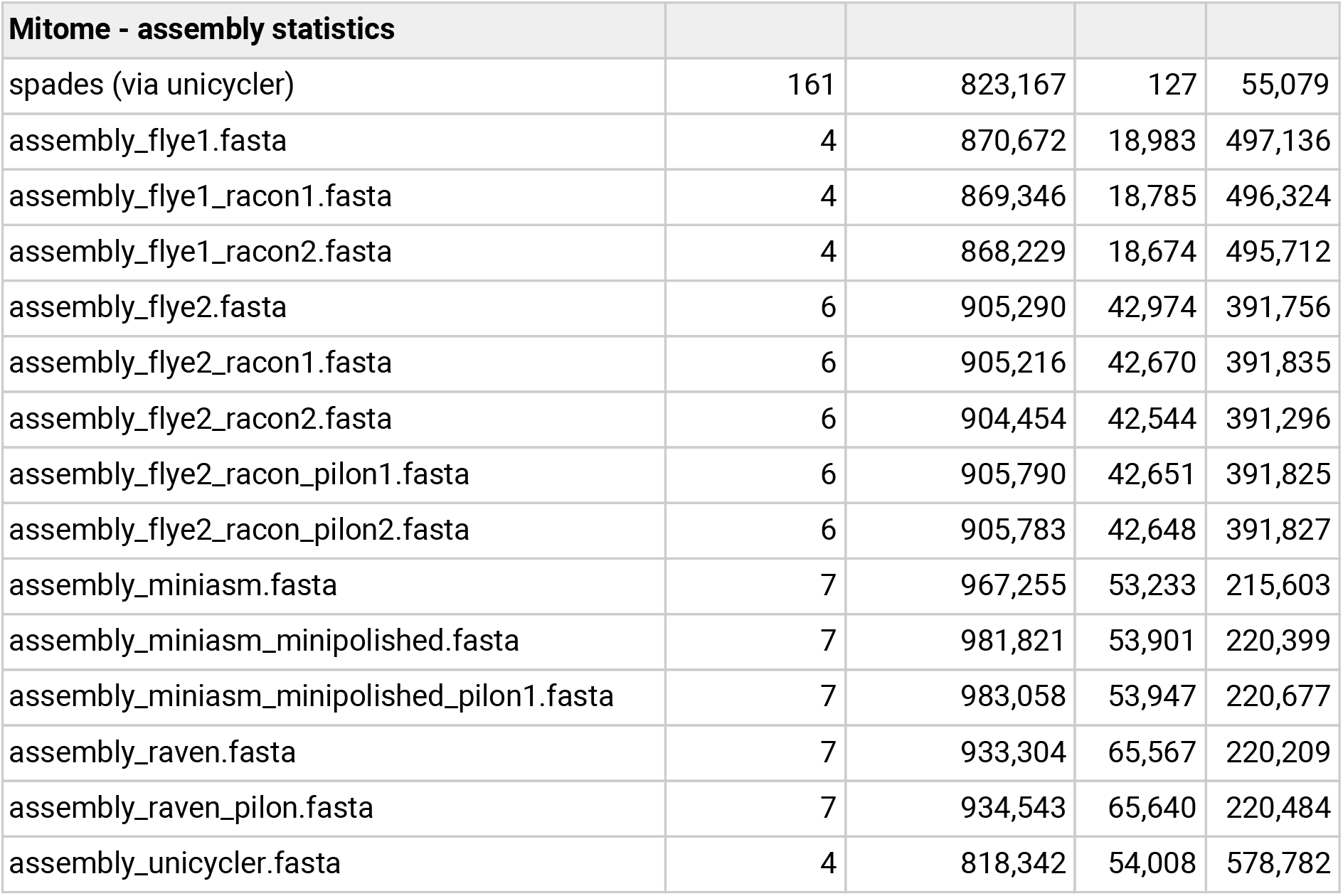
Read and assembly statistics.

### Read trimming and filtering

Nanopore reads were used raw with no read correction, as correction can introduce artificial consensus sequences, and because we have additional, more accurate data (Illumina) available for correction. To ensure Illumina reads were as accurate as possible for the correction step, we used fastp (S. Chen et al. 2018) to filter and trim reads, with settings: discard read or pair if more than 3 Ns; require min length 130; require average quality 35. This reduced number of read pairs to ~410 million (Table 1).

### Extraction and assembly of organelle-only Nanopore reads, round 1

To extract the organelle-only reads from the full read sets, we used a set of known sequences from related taxa as “baits”. For the plastome, we used three coding sequences from *Acacia ligulata* (NC_026134.2) in FASTA nucleotide format. We chose the genes *rbcL*, *matK* and *ndhF* as these are all likely to be plastid-only genes and are also well conserved. The *rbcL* and *matK* genes are usually located at either end of the LSC region, and *ndhF* is usually in the SSC region; these are well spaced around the plastid so that long reads should be extracted with roughly even coverage. As the mitome is much larger than the plastome, we used all 38 of the coding sequences from the mitome *Acacia ligulata* (NC_040998.1).

We mapped the raw Nanopore reads (~5.5 million) to the baits with minimap2 (Li 2018) and used samtools (Li et al. 2009) to extract mapped reads. We then used Filtlong (R. Wick n.d.) to keep only the longest of the extracted reads up to a coverage of X250, because assembly becomes more fragmented or not possible when coverage is too high (and preliminary tests confirmed this with our data). For the plastome, we extracted ~28,000 reads, downsampled to 901 reads, longest ~121 Kbp; for the mitome, we extracted ~14,000 reads, no downsampling as coverage did not meet cutoff (X250), longest ~105 Kbp (Table 1). Extracted Nanopore reads were assembled with Flye (Kolmogorov et al. 2019) and the assembly was polished with two rounds of Racon (Vaser et al. 2017).

### Extraction and assembly of organelle-only Nanopore reads, round 2

We used this first assembly as the baits file for the next round of extracting organelle reads from the original full read set. In Minimap2, we set a minimum match value to 5000, as preliminary tests showed that more leniency here resulted in too many reads extracted to assemble properly. Again we kept only the longest reads to a target coverage of X250. From the ~5.5 million raw reads, for the plastome, we extracted ~70,000 reads (approx twice as many as in round 1), downsampled to 864 reads, longest ~121,000 bp (same as round 1); for the mitome, we extracted ~14,000 reads (similar to round 1), downsampled slightly to ~12,000 reads, longest ~105 Kbp (same as round 1)(Table 1). As in the first round, these reads were then assembled with Flye and polished with two rounds of Racon. In testing, further rounds of Racon polishing made little difference.

### Extraction of organelle-only Illumina reads

Using the Round 2 assembly as baits, we then extracted organelle-only reads from the filtered and trimmed Illumina reads (~410 million read pairs). The extracted read sets were then randomly downsampled to a coverage of X250 using Rasusa (Hall 2020). For the plastome, this resulted in ~26 million read pairs, downsampled to ~130,000 read pairs; for the mitome, this resulted in ~23 million read pairs, downsampled to ~670,000 read pairs (Table 1).

### Polishing the assembly with Illumina reads

The round 2 assembly was then polished with the extracted, downsampled Illumina reads, using two rounds of Pilon (Li 2013; Walker et al. 2014), with a mindepth of 0.5 and fix set to bases (not contig breaks).

### Unicycler assembly

Using both the extracted Illumina and Nanopore reads, we used Unicycler (R. R. Wick et al. 2017) to perform a hybrid assembly. Unicycler first performs a short-read only assembly using Spades (Bankevich et al. 2012), and scaffolds this with long reads.

### Miniasm and Raven assemblies

Using the same read sets as used in Unicycler (long and short reads), further polished long-read assemblies were made. The Nanopore reads were assembled with Miniasm (Li 2016) + Minipolish (R. R. Wick and Holt 2019), and separately also with Raven (Vaser and Šikić 2020). Both assemblies were then further polished using Pilon (Li 2013; Walker et al. 2014) with the Illumina reads.

### Assembly comparisons and verification

Assembly graphs were visualized with the Bandage tool GUI (R. R. Wick et al. 2015). In particular, we used the BLAST (Altschul et al. 1990) tool within Bandage to compare assemblies. After loading a genome graph, a local BLAST database was built, and the query assembly file was compared; the assembly graph was then coloured by BLAST hits. We did several comparisons: comparing each assembly to the Unicycler assembly, and comparing the Unicycler assembly to three closely-related taxa from NCBI reference genomes. Further read mapping was done to verify that long reads spanned multiple alternate structures (see below).

### Scripts and computation

Computation details: GNU/Linux OS, 16 CPUs, 32GB RAM, 3TB disk. Custom script: assembler.sh, available at https://github.com/AnnaSyme/organelle-assembly, with initial baits files, Illumina adapters, and a conda yaml file (with tools and versions). Plastome parameters: input genome size of 160,000 (size only has to be approximate) and target bases (for filtering) of 40 Mb (= coverage X250). Mitome parameters: input genome size of 800,000 and target bases 200 Mb (= coverage X250).

### Draft annotation

We used the web service GeSeq v1.84 https://chlorobox.mpimp-golm.mpg.de/geseq.html to annotate the Unicycler assemblies, which primarily uses BLAT for sequence comparison, and as recommended (Tillich et al. 2017) used default settings but enabled tRNAScanSE and ARAGORN. Settings for both organelles were: enable BLAT search (Kent 2002) with Protein search identity: 25; rRNA, tRNA, DNA search identity: 85; ARAGORN v1.2.38 (Laslett and Canback 2004) with “Allow overlaps” and “Fix introns” enabled), tRNAscan-SE v2.0.6 (Chan et al. 2019), and OGDRAW v1.3.1 (Greiner, Lehwark, and Bock 2019) for visualisation.

Specific settings for the plastome were: an additional HMMER profile search (Wheeler and Eddy 2013) enabled with chloroplast land plants, ARAGORN with genetic code for plant chloroplast, MPI-MP chloroplast references enabled. Specific settings for mitome were: ARAGORN genetic code standard; BLAT reference sequence NCBI RefSeq *Acacia ligulata* mitome NC_040998.1.

The annotations are summarized into output GenBank and GFF3 files. No additional manual editing or curation was performed, so these annotations act only as a first-pass overview of gene and feature content of the assemblies.

## Results

### Overview

Plastome or mitome reads were extracted from the full read sets of Nanopore and Illumina data (Table 1). Short reads were assembled with Spades within Unicycler. Long reads were assembled with Flye, Miniasm and Raven, and assemblies were polished with long reads and then short reads. A hybrid assembly was performed with Unicycler. Assembly statistics are shown in Table 1, and summarised in Tables 2 and 3. The supplementary files include assemblies, assembly graphs, and annotation files; full details are listed at the end of the paper under “Code and data availability”.

**Table 2.** Plastome - main assembly results.

| **Assembly type** | **Tool** | **Total length** | **Contigs** | **Shortest contig** | **Longest contig** |
| --- | --- | --- | --- | --- | --- |
| short-read | spades | 135,939 | 12 | 128 | 62,974 |
| long-read | flye, polished | 173,695 | 2 | 80,947 | 92,748 |
| hybrid | unicycler | 173,902 | 1 | 173,902 | 173,902 |

**Table 3.** Mitome - main assembly results.

| **Assembly type** | **Tool** | **Total length** | **Contigs** | **Shortest contig** | **Longest contig** |
| --- | --- | --- | --- | --- | --- |
| short-read | spades | 823,167 | 161 | 127 | 55,079 |
| long-read | flye, polished | 905,783 | 6 | 42,648 | 391,827 |
| hybrid | unicycler | 818,342 | 4 | 54,008 | 578,782 |

### Plastome short-read assembly

Despite being based on short-reads only, the plastome assembly is fairly well-resolved: there are 12 contigs, smallest contig: 128 bp, largest contig: 62,974 bp, total length: 135,939 bp (Table 2, Figure 2). The assembly graph suggests the typical quadripartite structure of a long single-copy regions (LSC) as the larger circle in the graph, joined to inverted repeats (IRs) and a small single-copy region (SSC). The LSC has some unresolved repeats, as short reads (150 bp) cannot bridge these and place unambiguously in the assembly. The IR is a collapsed repeat of approximately double the coverage. The total size is shorter than the expected ~160 Kbp, because the inverted repeat is only counted once.

**Figure 2.**
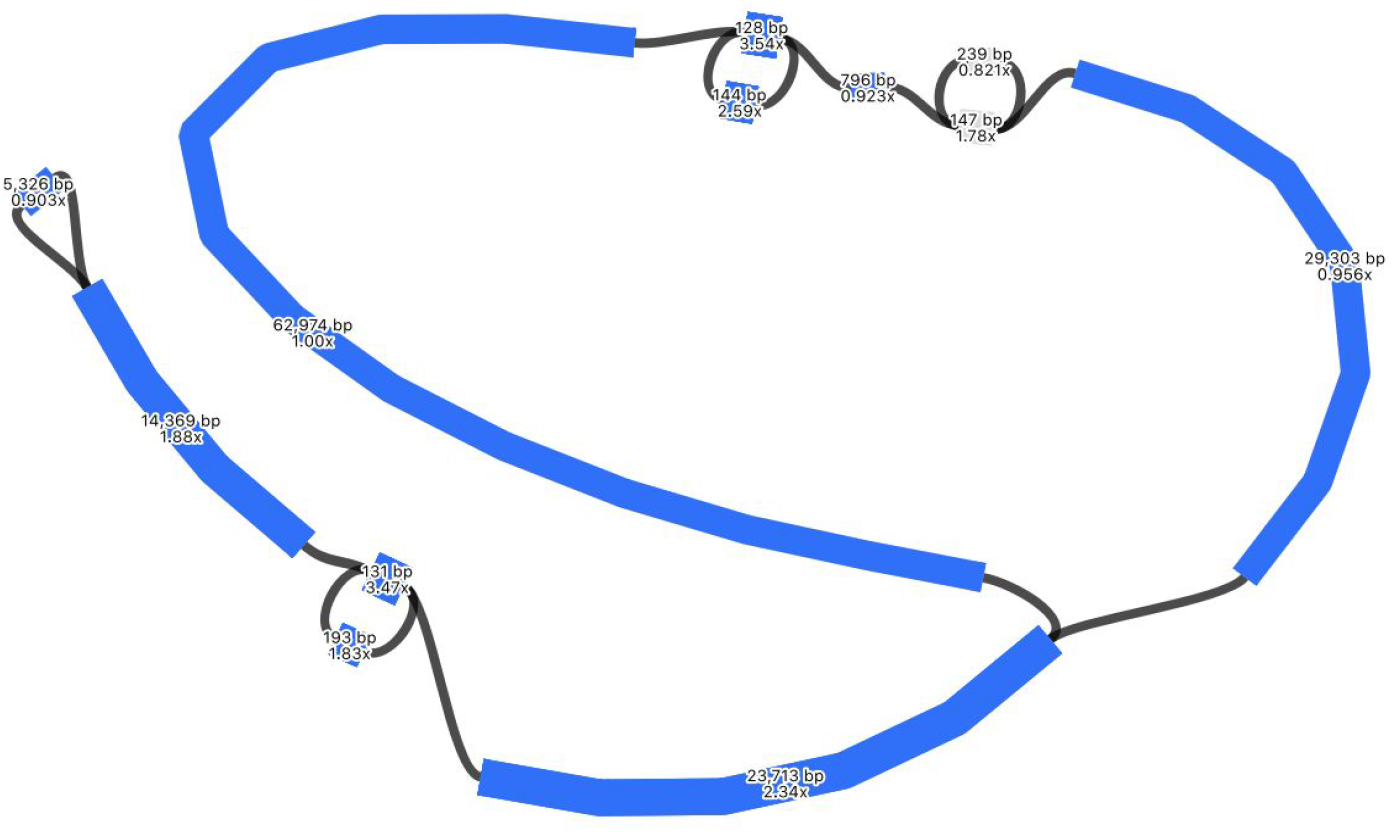
Assembly graph of the plastome based on short-read assembly with Spades, produced in the tool Bandage. Contigs are coloured according to their match with the hybrid Unicycler plastome assembly, using the BLAST tool within Bandage. Labels show contig lengths and depths.

### Plastome long-read assembly

As expected, this assembly is more fully resolved than the short-read assembly (Table 2, Figure 3). The only apparent collapsed repeat is the IR, separating the LSC and SSC. There are two contigs ~81 Kbp (IR + SSC + IR), ~92 Kbp (LSC). This assembly was then polished with the long reads (using Racon) and the short reads (using Pilon) which slightly increased the overall size by ~700 bp. The polished contig sizes are LSC (92,748 bp) and SSC joined by a collapsed IR (80,947 bp), total length: 173,695 bp (see Table 1 for all statistics).

**Figure 3.**
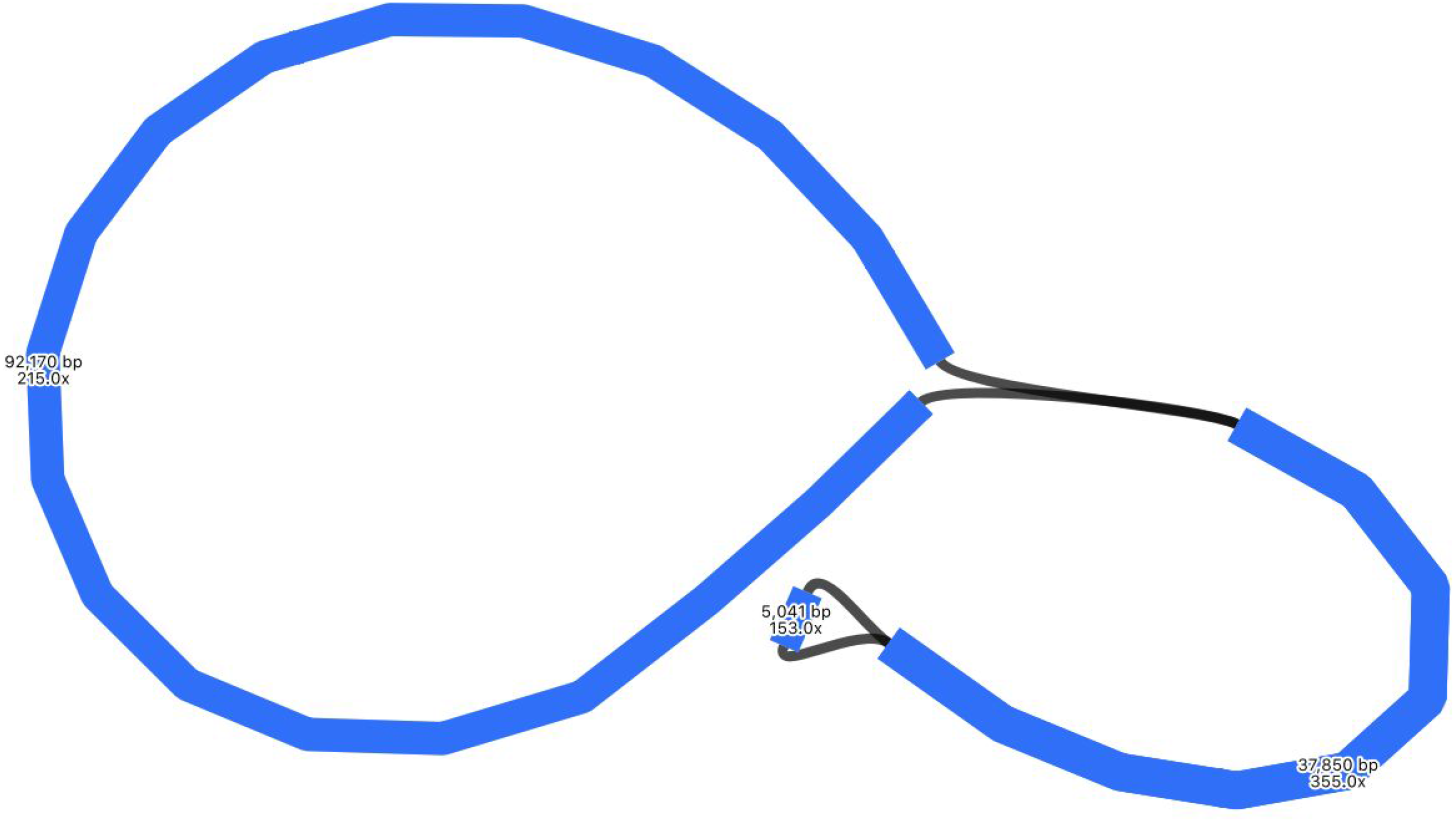
Assembly graph of the plastome based on a long-read assembly with Flye, produced in the tool Bandage. Contigs are coloured according to their match with the hybrid Unicycler plastome assembly, using the BLAST tool within Bandage. Labels show contig lengths and depths. This graph is unpolished; contig sizes differ slightly after polishing with both Racon and Pilon.

### Plastome hybrid assembly

The hybrid assembly by Unicycler resolved the plastome assembly into a single circle, of length 173,902 bp, which is very similar to the long-read polished assembly size 173,695 bp (Table 2, Figure 4). As Unicycler is designed to work well for hybrid read sets like this and can make use of the short read accuracy and long read bridging, we consider this the best representation of the plastome in this analysis. Although this assembly is resolved into a circle, we do keep in mind that there are likely two orientations of the SSC placement (Wang and Lanfear 2019) and suspect that the long-read assembly alone does not call consensus on this ambiguity.

**Figure 4.**
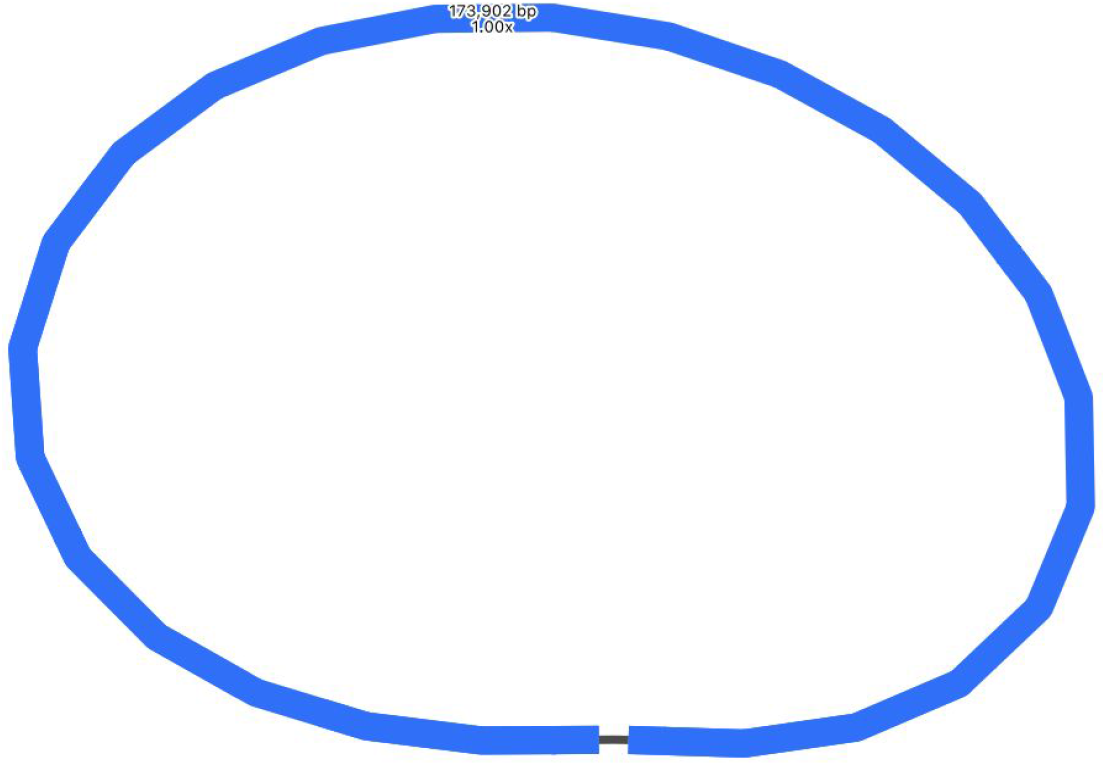
Assembly graph of the plastome based on a hybrid assembly with Unicycler, produced in the tool Bandage. Labels show contig length and depth.

### Plastome assemblies with other tools

Plastomes were also assembled using two other tools. Long reads were assembled with Miniasm, then polished with Minipolish and Pilon, producing an assembly of ~275 Kbp (Table 1, Figure 5). This assembly is much longer than both the Flye and Unicycler assemblies. Miniasm makes unitigs using the overlap-layout method, but with no consensus step. Here, either due to sequencing error and/or the multiple SSC orientations, it has likely assembled very similar regions which have not been collapsed into a consensus. Using BLAST, we can see that this is likely the case, as almost the entire Miniasm assembly matches the Unicycler assembly (Figure 5). To better visualise the components of this assembly, we used BLAST to find locations of the LSC, SSC and IRs, taken from the Flye assembly in Figure 3 (Figure 6). Here we can see that Miniasm has assembled reads into two contigs, one of which is almost the entire plastome, but that there is some ambiguity in how the other contig overlaps.

**Figure 5.**
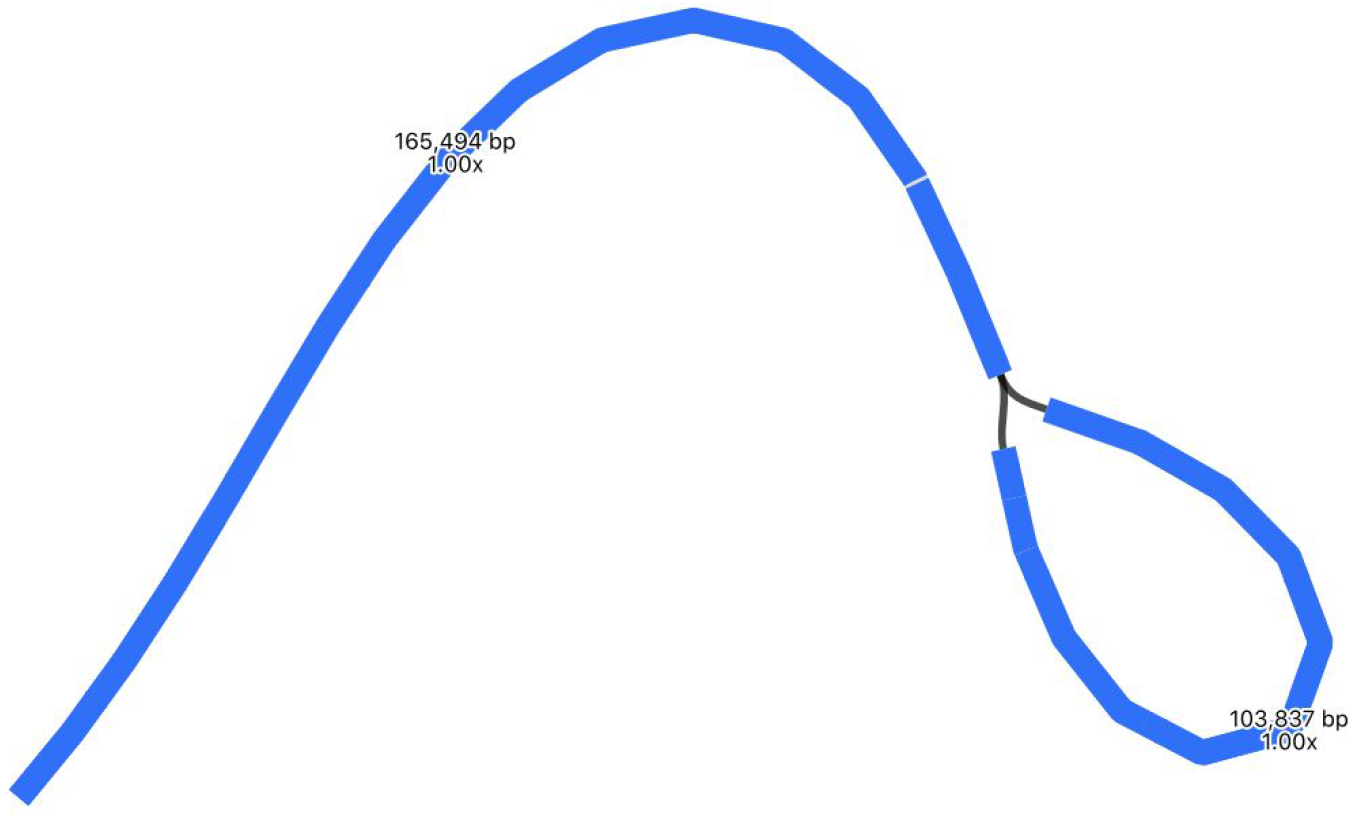
Assembly graph of the plastome based on a long-read assembly with Miniasm, produced in the tool Bandage. Contigs are coloured according to their match with the hybrid Unicycler plastome assembly, using the BLAST tool within Bandage. The whole assembly is covered showing that there is no new assembly here, only repeats of assembly sections that are present in the Unicycler assembly. Labels show contig lengths and depths. This graph is unpolished; contig sizes differ slightly after polishing with Pilon.

**Figure 6.**
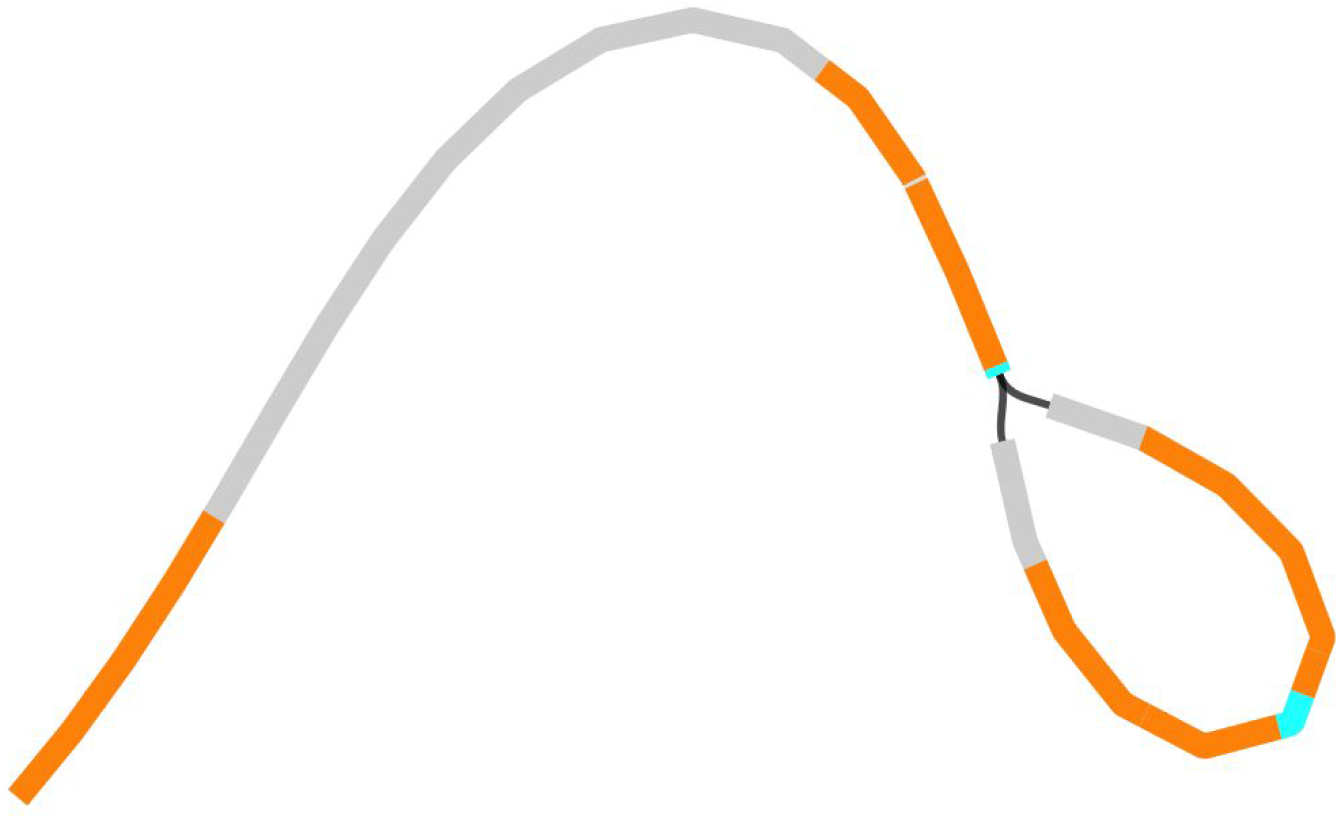
Assembly graph of the plastome based on a long-read assembly with Miniasm, produced in the tool Bandage. Contigs are coloured according to their match with particular sections of the Flye assembly: regions coloured grey match the LSC, regions coloured blue match the SSC, and regions in orange match the IR.

Long reads were also assembled with Raven, then polished with short reads and Pilon, producing a single contig of ~210 Kbp (Table 1, Figure 7). Raven uses OLC in a slightly different way to Miniasm, and then includes a consensus step using Racon. Thus, because it is using OLC, this assembly is longer than the Flye/Unicycler assemblies, but because it includes a consensus step, it is shorter than the Miniasm assembly. Again, to better visualize the components of this assembly, we used BLAST to compare it to the LSC, SSC and IR regions taken from the Flye assembly in Figure 3 (Figure 8) where we can see that the IR has been assembled approximately three times, and the SSC twice.

**Figure 7.**
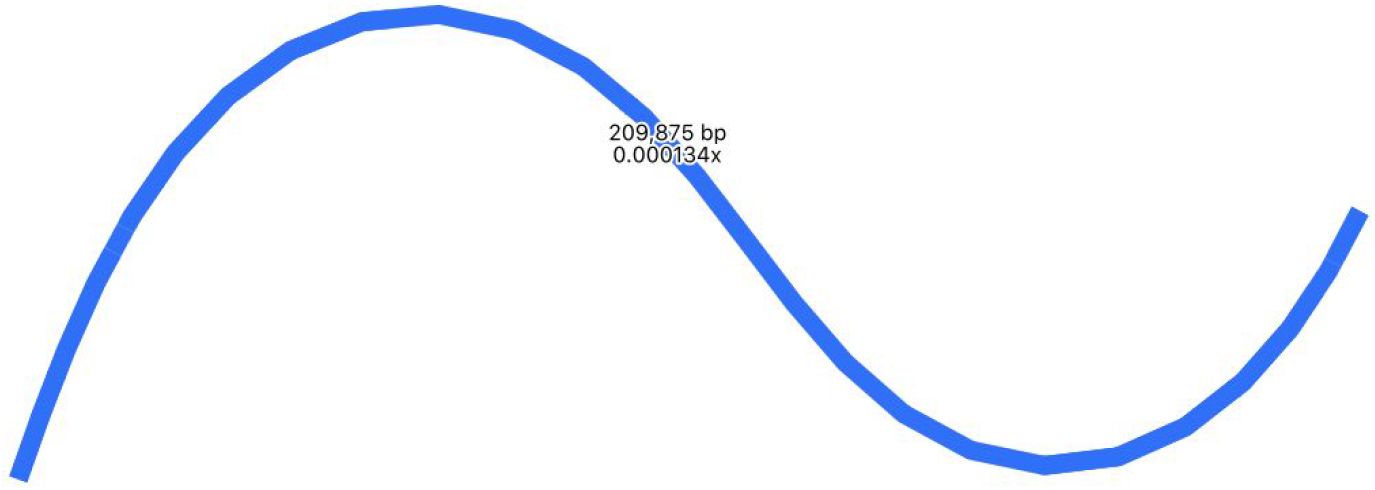
Assembly graph of the plastome based on a long-read assembly with Raven, produced in the tool Bandage. Contigs are coloured according to their match with the hybrid Unicycler plastome assembly, using the BLAST tool within Bandage. The whole assembly is covered showing that there is no new assembly here, only repeats of assembly sections that are present in the Unicycler assembly. Labels show contig lengths and depths. This graph is unpolished; contig sizes differ slightly after polishing with Pilon.

**Figure 8.**
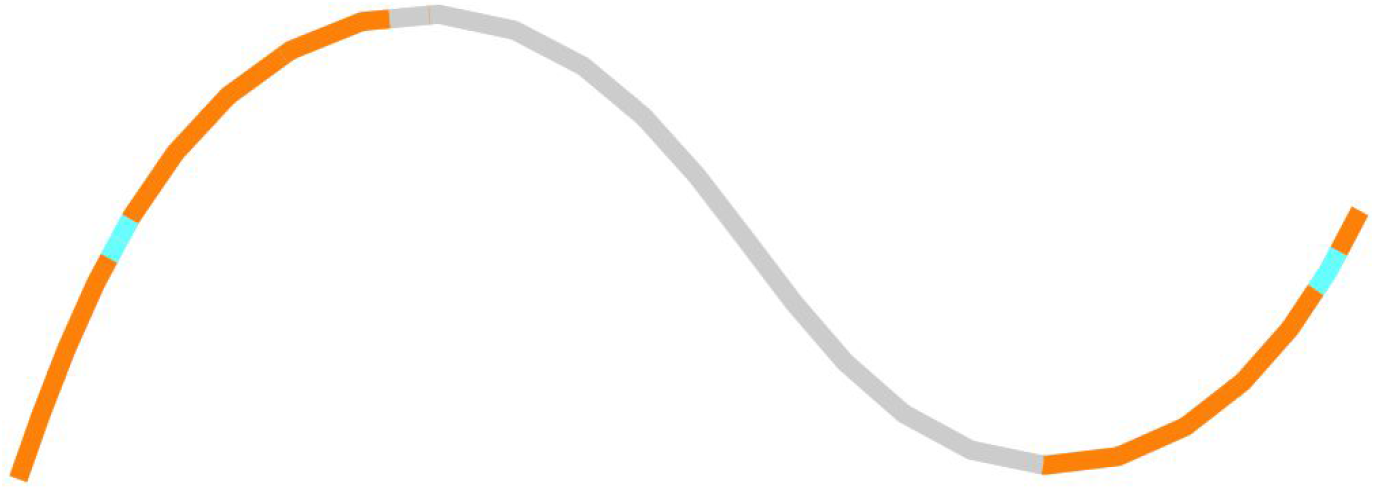
Assembly graph of the plastome based on a long-read assembly with Miniasm, produced in the tool Bandage. Contigs are coloured according to their match with particular sections of the Flye assembly: regions coloured grey match the LSC, regions coloured blue match the SSC, and regions in orange match the IR.

### Plastome draft annotation

The draft annotation (Figure 9) is a visual first-pass approximation of the gene and feature content rather than a highly polished finished annotation. The supplementary files including GenBank and GFF3 formats of this annotation are available for researchers to further explore this annotation (see “Code and data availability”).

**Figure 9.**
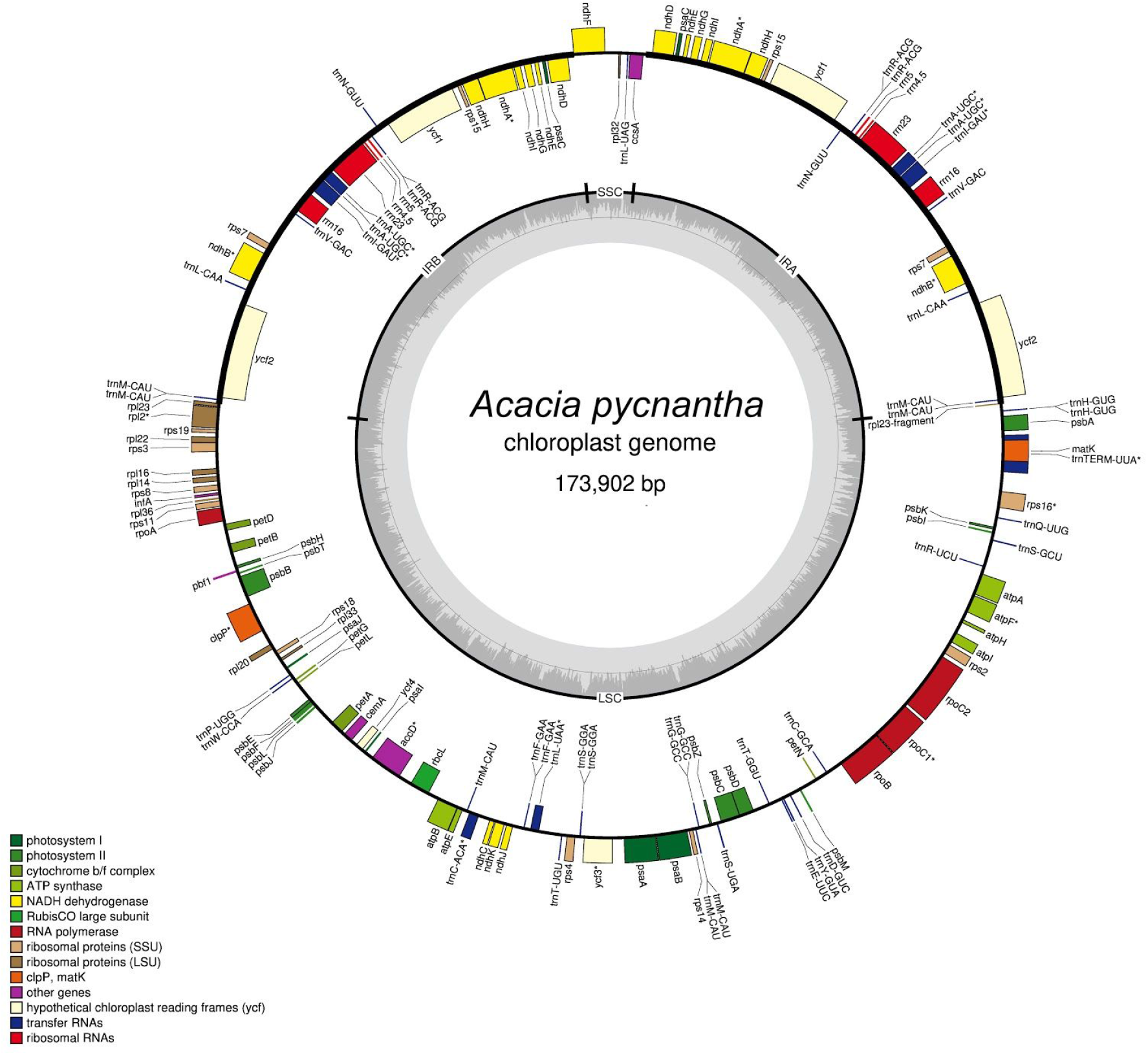
Annotated plastome of Acacia pycnantha, based on Unicycler assembly, and produced by the GeSeq tool and OGDRAW.

### Plastome summary

Based on the Unicycler assembly here and annotations by GeSeq, the assembly of the *Acacia pycnantha* plastome is broadly similar to previous results found in other *Acacia* species (Table 4). As an additional visual comparison, we used the BLAST tool within Bandage to compare the Unicycler assembly of *Acacia pycnantha* with plastomes of related species in subfamily Caesalpinioideae: *Acacia ligulata* (NCBI Reference Sequence NC_026134.2), *Leucaena trichandra* (NCBI Reference Sequence NC_028733.1), and *Haematoxylum brasiletto* (GenBank KJ468097.1). There are no large unmatched sections in the *Acacia pycnantha* assembly that would indicate potentially novel regions or misassembly (Figure 10).

**Figure 10.**
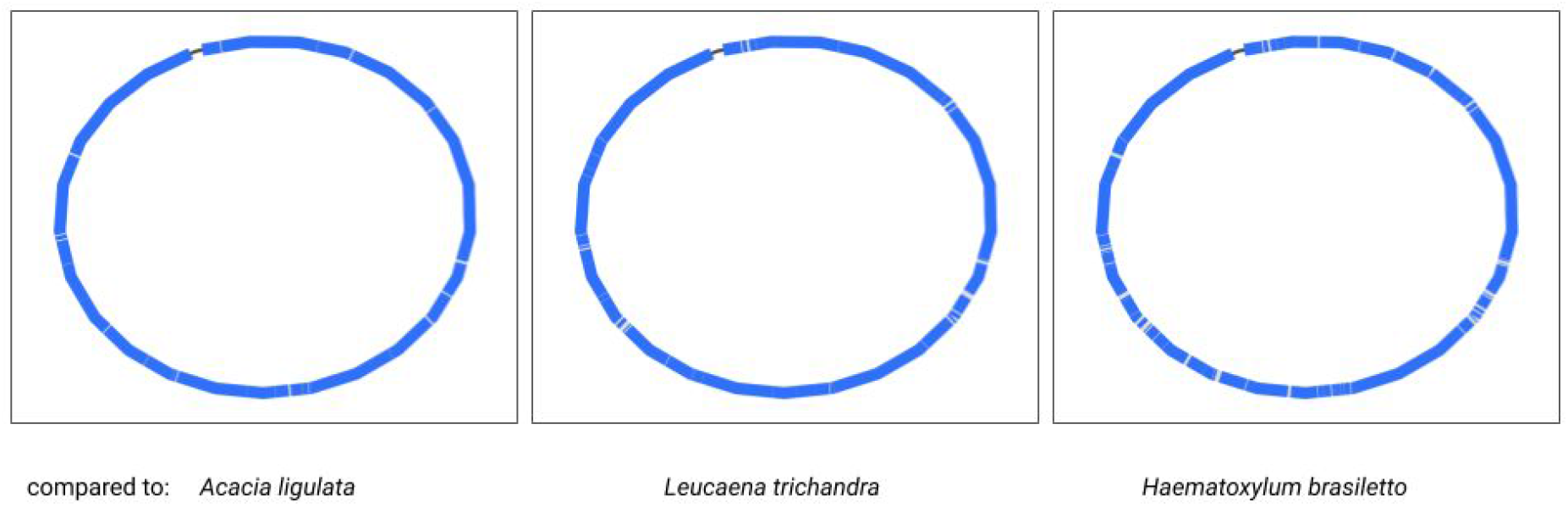
Comparison of the Acacia pycnantha Unicycler assembly with plastomes from related species. The contig is coloured according to its match with these assemblies, using the BLAST tool within Bandage. No novel regions or misassembly are evident within Acacia pycnantha.

**Table 4.**
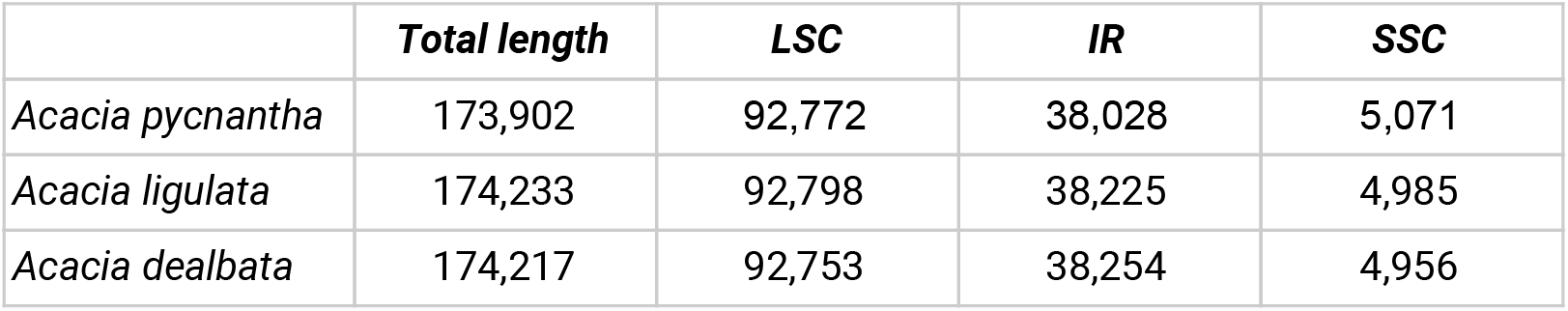
Comparisons between different Acacia plastome assemblies, showing number of base pairs in different genome components. Acacia ligulata statistics are from (Williams et al. 2015); Acacia dealbata statistics are from (Asaf et al. 2019). Acacia pycnantha statistics are derived from the GeSeq annotation, visualized in OGDRAW.

|  | **Total length** | **LSC** | **IR** | **SSC** |
| --- | --- | --- | --- | --- |
| *Acacia pycnantha* | 173,902 | 92,772 | 38,028 | 5,071 |
| *Acacia ligulata* | 174,233 | 92,798 | 38,225 | 4,985 |
| *Acacia dealbata* | 174,217 | 92,753 | 38,254 | 4,956 |

### Mitome short-read assembly

The mitome assembly based on short reads has 161 contigs, ranging from 127 bp to ~55,000 bp in length, to give a total size of 823,167 bp (Table 3). As expected, the assembly graph shows a fair amount of unresolved ambiguity, at least one dead end, and several very small fragments (Figure 11).

**Figure 11.**
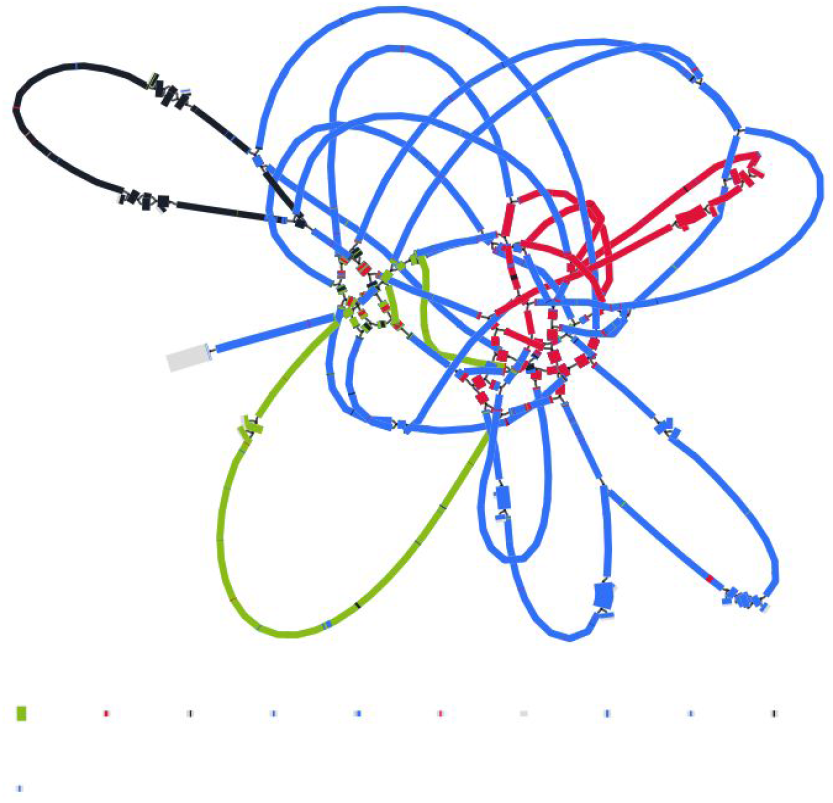
Assembly graph of the mitome based on short-read assembly with Spades, produced in the tool Bandage. Contigs are coloured according to their match with the hybrid Unicycler mitome assembly, using the BLAST tool within Bandage.

### Mitome long-read assembly

The long-read assembly of the mitome is a vast improvement over the short-read assembly (Table 3) in terms of contig lengths and contiguity. The number of contigs has reduced from 161 to 6, the shortest contig has increased in size from 127 bp to ~43 Kbp, and the longest has increased from ~55 Kbp to ~392 Kbp. Total length has increased from ~823 Kbp to ~906 Kbp. The assembly graph is much less tangled: there are two possibly circular segments of ~93 Kbp and ~108 Kbp, and the remainder forms a single structure albeit with some ambiguous regions (Figure 12). A note that contigs in the assembly graph are different to the contigs in the FASTA file: FASTA file contigs include only the longest unambiguous paths and so are broken at repeats.

**Figure 12.**
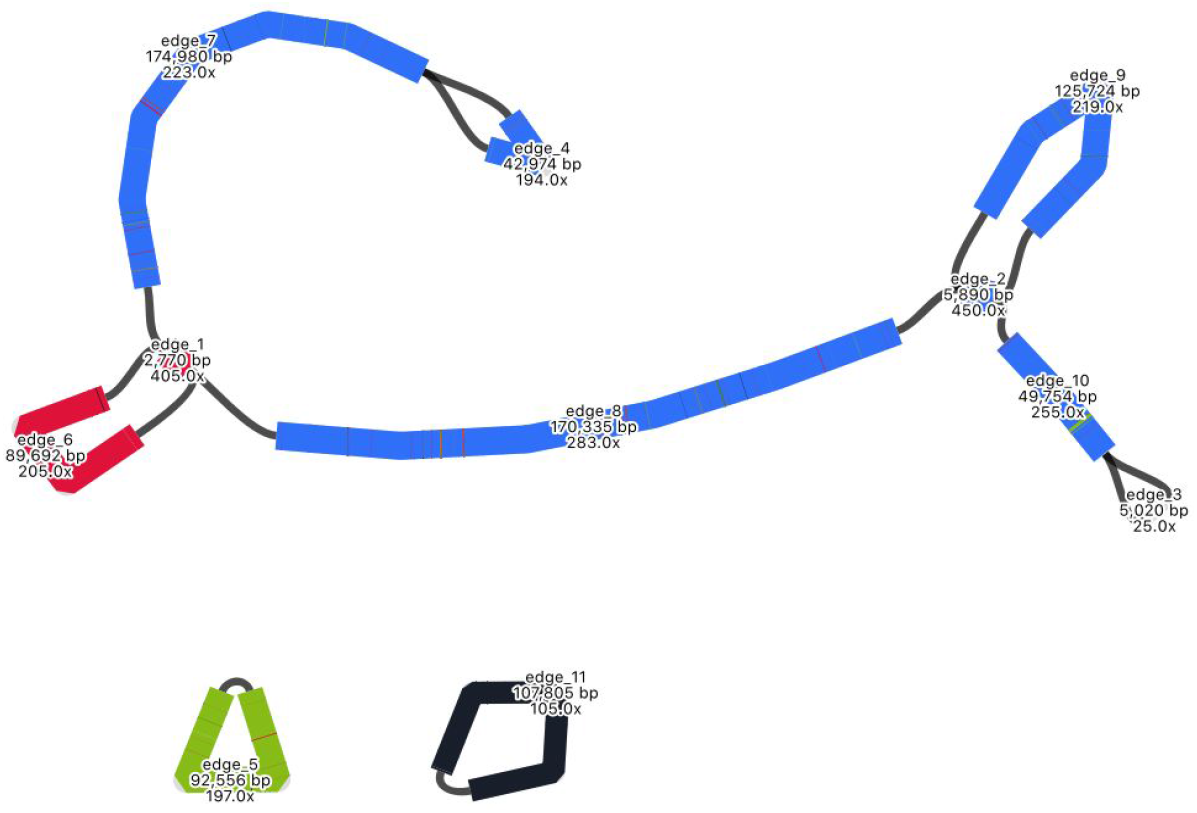
Assembly graph of the mitome based on long-read assembly with Flye, produced in the tool Bandage. Contigs are coloured according to their match with the hybrid Unicycler mitome assembly, using the BLAST tool within Bandage. Labels show contig lengths and depths. This graph is unpolished; contig sizes differ slightly after polishing with both Racon and Pilon.

### Mitome hybrid assembly

The mitome hybrid assembly produced by Unicycler is well resolved, with some apparent improvements over the long-read assembly by Flye (Table 3, Figure 13). The number of contigs has decreased from 6 to 4, shortest contig has increased in size from ~43 Kbp to ~54 Kbp, and the longest contig from ~392 Kbp to ~579 Kbp. Total size has decreased from ~906 Kbp to ~818 Kbp. There are three apparent circular segments of sizes ~93 Kbp, ~93 Kbp and ~54 Kbp.

**Figure 13.**
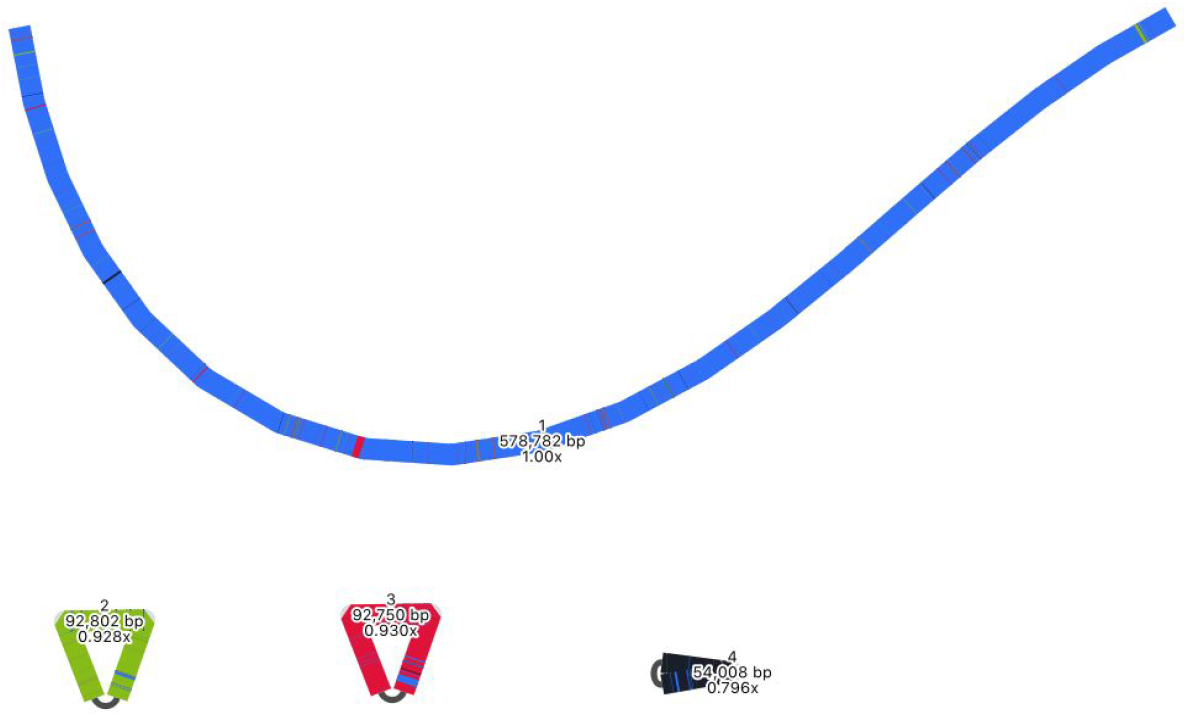
Assembly graph of the mitome based on a hybrid assembly with Unicycler, produced in the tool Bandage. Labels show contig names, length and depth. To compare this assembly with the other assemblies in this analysis, contigs have been coloured according to their match with self, using the BLAST tool within Bandage.

As with the plastome assembly, based on the strengths of Unicycler in working with hybrid read sets, we consider this Unicycler assembly the best representation of the mitome in this analysis. However, by also considering the long-read Flye assembly, we can explore the complexity of this genome structure further. The Flye assembly joins a longer section together, indicating how a particular segment - coloured in red - may be integrated. Although the Flye assembly is substantially longer than the Unicycler assembly, a BLAST comparison (Figure 12) shows that all components match to the Unicycler assembly well, indicating that additional length may be from a similar repeat region that has not been collapsed. The Flye assembly also shows that one of its circular segments - coloured in black - is twice the size of the similar segment in the Unicycler assembly, which again suggests a repeat region that has not been collapsed. Whether these repeat regions are truly independent and should be collapsed is unclear, and demonstrates that the complexity of this structure is not trivial to resolve.

### Mitome assemblies with other tools

Mitomes were also assembled with two other tools. Long reads were assembled with Miniasm, then polished with Minipolish and Pilon, producing an assembly of ~983 Kbp (Table 1, Figure 14). As with the plastome results, this assembly is much longer than the Flye and Unicycler assemblies, which is expected as Miniasm has no consensus step. Using BLAST to compare this assembly to that produced by Unicycler, we can see that all sections match the Unicycler assembly (Figure 14).

**Figure 14.**
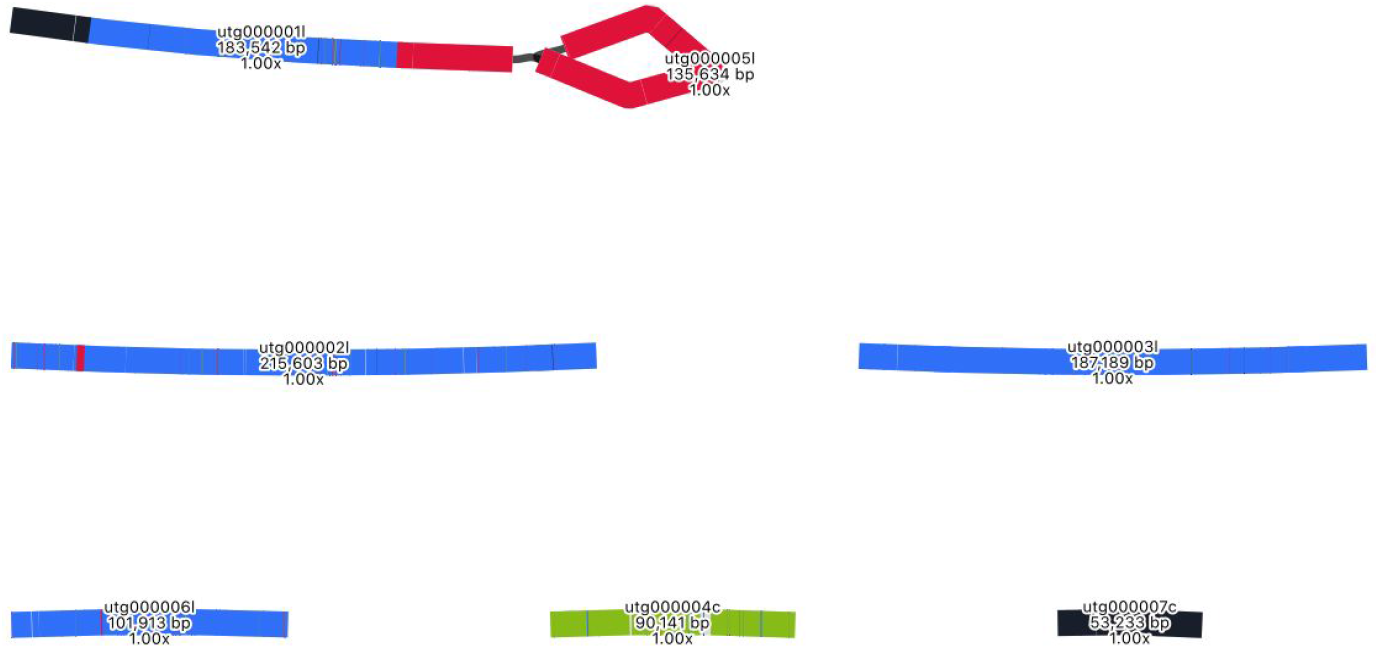
Assembly graph of the mitome based on a long-read assembly with Miniasm, produced in the tool Bandage. Contigs are coloured according to their match with the hybrid Unicycler mitome assembly, using the BLAST tool within Bandage. The whole assembly is covered showing that there is no new assembly here, only repeats of assembly sections that are present in the Unicycler assembly. Labels show contig names, lengths and depths. This graph is unpolished; contig sizes differ slightly after polishing with Pilon.

Long reads were also assembled with Raven, then polished with short reads and Pilon, producing an assembly of ~935 Kbp (Table 1, Figure 15). As with the plastome results, this Raven assembly is shorter than the Miniasm assembly, but longer than the Flye/Unicycler assemblies, due to the algorithm employed. A BLAST comparison with the Unicycler assembly confirms that all sections match (Figure 15).

**Figure 15.**
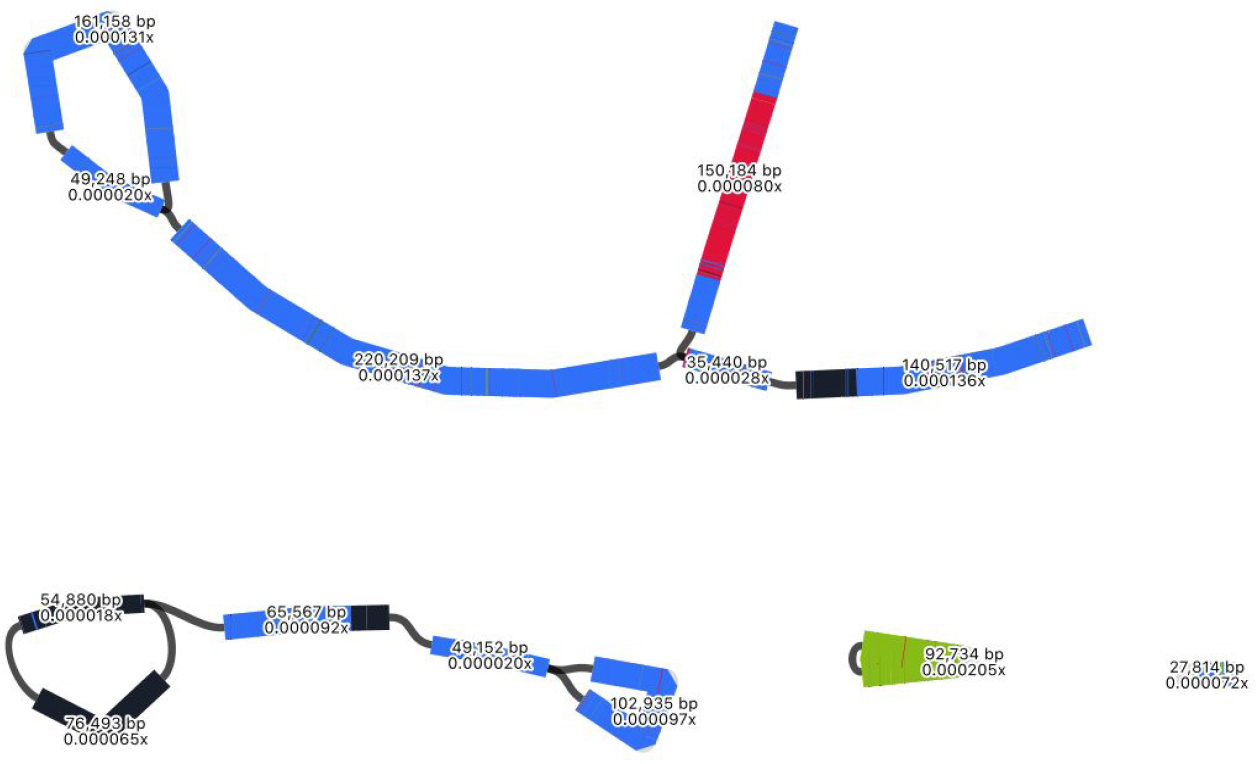
Assembly graph of the mitome based on a long-read assembly with Raven, produced in the tool Bandage. Contigs are coloured according to their match with the hybrid Unicycler mitome assembly, using the BLAST tool within Bandage. The whole assembly is covered showing that there is no new assembly here, only repeats of assembly sections that are present in the Unicycler assembly. Labels show contig lengths and depths. This graph is unpolished; contig sizes differ slightly after polishing with Pilon.

An interesting result from the Minisam and Raven assemblies is the placement of a particular segment, coloured in black. In the Unicycler assembly, this segment is circularised, but in these assemblies there is an indication of how the segment may join to other parts of the genome (Figures 14, 15).

### Mitome draft annotation

The draft annotation (Figure 16) is presented to provide a first-pass visualisation of gene and feature content. To further explore this annotation, supplementary files include GenBank and GFF3 formats of this annotation (see “Code and data availability”).

**Figure 16.**
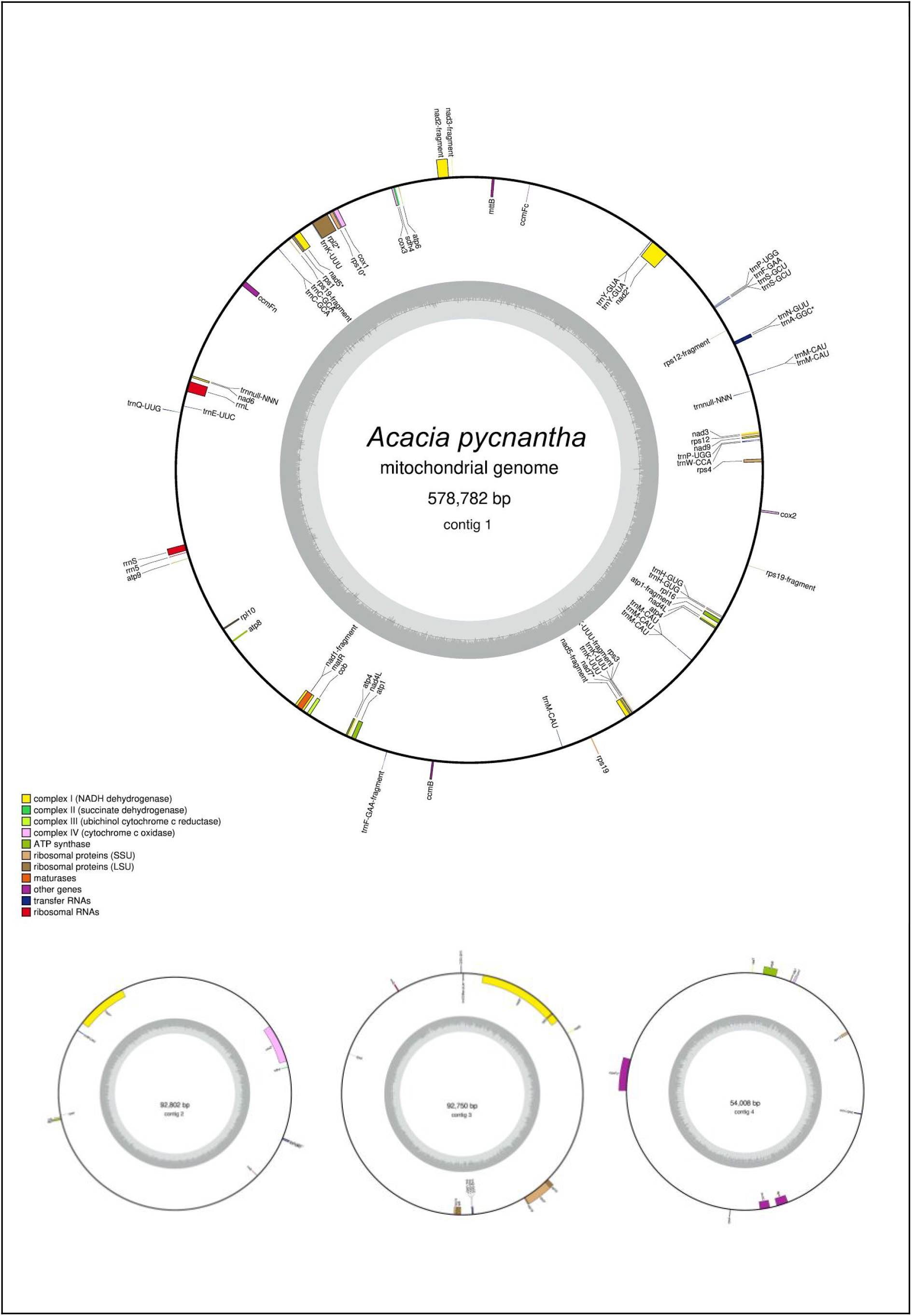
Annotated plastome of Acacia pycnantha, based on Unicycler assembly, and produced by the GeSeq tool and OGDRAW.

### Mitome summary

Based on the Unicycler assembly, the assembly of the *Acacia pycnantha* mitome is 818,342 bp, and may be arranged in a long linear piece and three smaller circular segments. Alternative assembly results suggest that some circular segments can be incorporated into the larger structure. In comparison, the closest sequenced relative, *Acacia ligulata* is substantially shorter, with a total length of 698,138 bp (Sanchez-Puerta et al. 2019). This was assembled with short Illumina reads into 10 contigs, followed by manual editing and joining. Interestingly, the work on *Acacia ligulata* suggested the possible existence of alternative structures in the form of head-to-tail concatemers, which is consistent with the alternate forms assembled for *Acacia pycnantha* mitomes herein.

As an additional visual comparison, with used the BLAST tool within Bandage to compare the Unicycler assembly of *Acacia pycnantha* with mitomes of related species in subfamily Caesalpinioideae: *Acacia ligulata* (NCBI Reference Sequence NC_040998.1), *Leucaena trichandra* (NCBI Reference Sequence NC_039738.1), and *Haematoxylum brasiletto* (NCBI Reference Sequence NC_045040.1). In contrast to the plastome assembly comparisons, there is a fair amount of non-matching sequence in the *Acacia pycnantha* assemblies, increasing in concert with phylogenetic distance (Figure 17).

**Figure 17.**
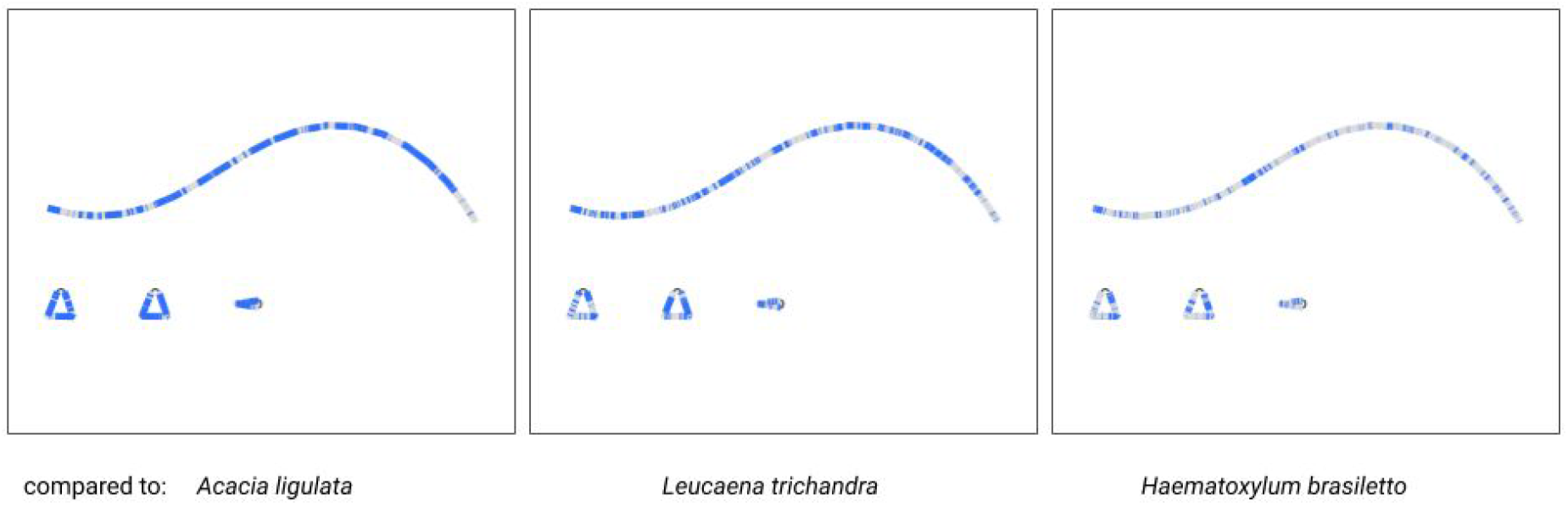
Comparison of the Acacia pycnantha Unicycler assembly with mitomes from related species. The contig is coloured according to its match with these assemblies, using the BLAST tool within Bandage. Increasing difference is seen in concert with increasing phylogenetic distance, from left to right.

### Mitome structural variation

One of the major aims of this paper is to understand more about multiple structures of an organelle genome that may exist simultaneously. Although the various results from assemblies do suggest this, we did an extra manual check of whether long reads would span multiple structures. To do this, we specifically looked at the location of the red contig, with a size of ~90 Kbp. In the long-read Flye assembly (Figure 12), this contig (= edge 6) is integrated into the large blue connected component, and located between the repeat region of 2,770 bp (= edge 1). However, in the Unicycler assembly (Figure 13), this red contig is excised as an independent circular contig, and a small fragment is also present in the long linear contig.

The question is: can both of these structures exist? Do long reads support both configurations? The two paths to compare in Figure 12 are a structure that includes edge 6 (edge 8 - edge 1 - edge 6 - edge 1 - edge 7) and a structure that excludes edge 6 (edge 8 - edge 1 - edge 7). We mapped the long reads to these paths to visually inspect whether there was support for both of these assemblies. To do this, we drew the Flye assembly in Bandage in double mode (Figure 18), extracted nodes in the two paths of interest (keeping direction consistent), reduced the lengths of the outer contigs (edges 7 and 8) to only 10,000 bp, and combined them into a single FASTA file of paths. Then we used this FASTA file as bait to extract matching reads from the full mitome Nanopore read set with minimap2 (Li 2018), and visualized the bam track in JBrowse (Buels et al. 2016) in Galaxy (Afgan et al. 2018)(See “Code and data availability” for bam file). This confirmed that long reads span both paths and thus both structural configurations, where the ~90 Kbp red contig can be incorporated into the larger mitome structure (Figure 12) or not (Figure 13).

**Figure 18.**
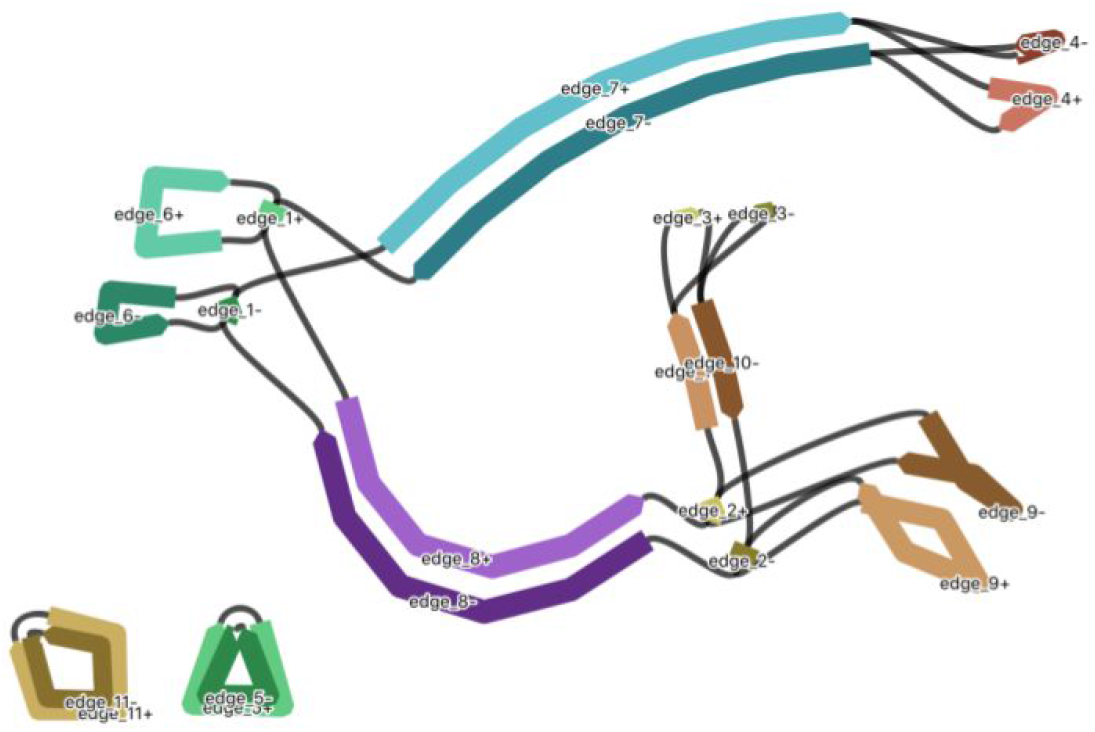
Tracing alternate paths through the mitome assembly graph, to include or exclude edge 6. Graph is drawn in Bandage in double mode, so the path directionality is maintained when nodes are extracted.

## Discussion

Long sequencing reads are becoming the new normal for genome assembly projects, providing new ways to investigate structural complexity. In this work, we were able to successfully extract organelle-only reads from full nuclear+organelle read sets, and assemble the reads under a variety of algorithms in well-tested tools. In this case, we consider the best representations of the *Acacia pycnantha* mitome and plastome to have been achieved by the hybrid short and long read assembly with Unicycler, and draft annotations have been presented from the GeSeq tool.

Additional assemblies have suggested the existence of multiple mitome configurations, a hypothesis supported by long reads that span alternate assembly graph paths. This builds on a body of work that increasingly refutes the existence of a single, static circular mitochondrial genome (Sloan 2013; Kozik et al. 2019; Wu et al. 2020; Jackman et al. 2020).

As there are many avenues to explore to improve both assemblies and annotations, we consider the assemblies presented here to represent version 1. New technologies are improving long read fidelity, such as PacBio HiFi sequencing. Oxford Nanopore raw sequencing data can benefit from being re base-called with new tools (Xu et al. 2020), and trained on relevant taxonomic data (R. R. Wick, Judd, and Holt 2019). Long-read specific assemblers are continually optimised, particularly to error profiles, and there is a large focus on improving the assembly of repeat regions (Bankevich and Pevzner 2020).

One option to explore in further research is that multiple structures may be better assembled via metagenomic approaches. Alternate structures could be considered part of a metagenomic pool, and reads clustered and assembled accordingly, taking into account that abundances of alternate forms may not be equal.

In either case, the increased use of long reads and *de novo* assembly will further improve organelle assemblies and pave the way for fuller genomic comparison across species.

## Abbreviations

bp: base pair
SSC: short single copy region
LSC: long single copy region
IR: inverted repeat
plastome: chloroplast genome
mitome: mitochondrial genome

## Code and data availability

Raw sequence data: Raw data will be made available in a public repository on publication. Supplementary files available at Zenodo (https://doi.org/10.5281/zenodo.4330088): for each organelle genome, these are 14 assemblies in fasta format, and associated GFA format file if available (not all stages produce this file), as well as the Spades GFA from Unicycler. For each Unicycler assembly there is a set of annotation files that include GenBank and GFF3 formats, and outputs from HMMER, ARAGORN, and tRNAscan-SE. There is also a bam file of reads mapped to alternate assembly paths for the mitome to explore the 90 Kbp contig of interest. A copy of the assembly script (assembler.sh) is available in Zenodo and at this repository - https://github.com/AnnaSyme/organelle-assembly - with instructions on how to run the script and the required inputs and tools.

## Acknowledgements

We would like to acknowledge the contribution of the Genomics for Australian Plants Framework Initiative consortium (https://www.genomicsforaustralianplants.com/consortium/) in the generation of data used in this publication. The Initiative is supported by funding from Bioplatforms Australia (enabled by NCRIS), the Ian Potter Foundation, Royal Botanic Gardens Victoria, the Royal Botanic Gardens and Domain Trust, CSIRO, Centre for Australian National Biodiversity Research and the Department of Biodiversity, Conservation and Attractions, Western Australia. Anna Syme wishes to thank Torsten Seemann for discussion on the iterative bait + assembly approach, and Shaun Jackman for discussion on mitochondrial genome structure and for suggesting the utility of the Blast tool within the Bandage assembly viewer.

## References

Afgan, Enis, Dannon Baker, Bérénice Batut, Marius van den Beek, Dave Bouvier, Martin Čech, John Chilton, et al. 2018. “The Galaxy Platform for Accessible, Reproducible and Collaborative Biomedical Analyses: 2018 Update.” Nucleic Acids Research 46 (W1): W537–44. https://doi.org/10.1093/nar/gky379.

Altschul, Stephen F., Warren Gish, Webb Miller, Eugene W. Myers, and David J. Lipman. 1990. “Basic Local Alignment Search Tool.” Journal of Molecular Biology 215 (3): 403–10. https://doi.org/10.1016/S0022-2836(05)80360-2.

Asaf, Sajjad, Arif Khan, Abdul Latif Khan, Ahmed Al-Harrasi, and Ahmed Al-Rawahi. 2019. “Complete Chloroplast Genomes of Vachellia Nilotica and Senegalia Senegal: Comparative Genomics and Phylogenomic Placement in a New Generic System.” PloS One 14 (11): e0225469. https://doi.org/10.1371/journal.pone.0225469.

Bankevich, Anton, Sergey Nurk, Dmitry Antipov, Alexey A. Gurevich, Mikhail Dvorkin, Alexander S. Kulikov, Valery M. Lesin, et al. 2012. “SPAdes: A New Genome Assembly Algorithm and Its Applications to Single-Cell Sequencing.” Journal of Computational Biology: A Journal of Computational Molecular Cell Biology 19 (5): 455–77. https://doi.org/10.1089/cmb.2012.0021.

Bankevich, Anton, and Pavel Pevzner. 2020. “mosaicFlye: Resolving Long Mosaic Repeats Using Long Error-Prone Reads.” https://doi.org/10.1101/2020.01.15.908285.

Bendich, A. J. 1996. “Structural Analysis of Mitochondrial DNA Molecules from Fungi and Plants Using Moving Pictures and Pulsed-Field Gel Electrophoresis.” Journal of Molecular Biology 255 (4): 564–88. https://doi.org/10.1006/jmbi.1996.0048.

Buels, Robert, Eric Yao, Colin M. Diesh, Richard D. Hayes, Monica Munoz-Torres, Gregg Helt, David M. Goodstein, et al. 2016. “JBrowse: A Dynamic Web Platform for Genome Visualization and Analysis.” Genome Biology 17 (1): 66. https://doi.org/10.1186/s13059-016-0924-1.

Chan, Patricia P., Brian Y. Lin, Allysia J. Mak, and Todd M. Lowe. 2019. “tRNAscan-SE 2.0: Improved Detection and Functional Classification of Transfer RNA Genes.” https://doi.org/10.1101/614032.

Chen, Shifu, Yanqing Zhou, Yaru Chen, and Jia Gu. 2018. “Fastp: An Ultra-Fast All-in-One FASTQ Preprocessor.” Bioinformatics 34 (17): i884–90. https://doi.org/10.1093/bioinformatics/bty560.

Chen, Zhiwen, Nan Zhao, Shuangshuang Li, Corrinne E. Grover, Hushuai Nie, Jonathan F. Wendel, and Jinping Hua. 2017. “Plant Mitochondrial Genome Evolution and Cytoplasmic Male Sterility.” Critical Reviews in Plant Sciences 36 (1): 55–69. https://doi.org/10.1080/07352689.2017.1327762.

Dierckxsens, Nicolas, Patrick Mardulyn, and Guillaume Smits. 2017. “NOVOPlasty: De Novo Assembly of Organelle Genomes from Whole Genome Data.” Nucleic Acids Research 45 (4): e18. https://doi.org/10.1093/nar/gkw955.

Frommer, Bianca, Daniela Holtgräwe, Ludger Hausmann, Prisca Viehöver, Bruno Huettel, Reinhard Töpfer, and Bernd Weisshaar. 2020. “Genome Sequences of Both Organelles of the Grapevine Rootstock Cultivar ‘Börner.’” Microbiology Resource Announcements 9 (15). https://doi.org/10.1128/MRA.01471-19.

Greiner, Stephan, Pascal Lehwark, and Ralph Bock. 2019. “OrganellarGenomeDRAW (OGDRAW) Version 1.3.1: Expanded Toolkit for the Graphical Visualization of Organellar Genomes.” Nucleic Acids Research 47 (W1): W59–64. https://doi.org/10.1093/nar/gkz238.

Guyeux, Christophe, Jean-Claude Charr, Hue T. M. Tran, Agnelo Furtado, Robert J. Henry, Dominique Crouzillat, Romain Guyot, and Perla Hamon. 2019. “Evaluation of Chloroplast Genome Annotation Tools and Application to Analysis of the Evolution of Coffee Species.” PloS One 14 (6): e0216347. https://doi.org/10.1371/journal.pone.0216347.

Hall, Michael. 2020. “Rasusa: Randomly Subsample Sequencing Reads to a Specified Coverage.” March 27, 2020. https://doi.org/10.5281/zenodo.3731394.

Jackman, Shaun D., Lauren Coombe, René L. Warren, Heather Kirk, Eva Trinh, Tina MacLeod, Stephen Pleasance, et al. 2020. “Complete Mitochondrial Genome of a Gymnosperm, Sitka Spruce (Picea Sitchensis), Indicates a Complex Physical Structure.” Genome Biology and Evolution, May. https://doi.org/10.1093/gbe/evaa108.

Jin, Jian-Jun, Wen-Bin Yu, Jun-Bo Yang, Yu Song, Claude W. dePamphilis, Ting-Shuang Yi, and De-Zhu Li. 2019. “GetOrganelle: A Fast and Versatile Toolkit for Accurate de Novo Assembly of Organelle Genomes.” https://doi.org/10.1101/256479.

Kent, W. James. 2002. “BLAT--the BLAST-like Alignment Tool.” Genome Research 12 (4): 656–64. https://doi.org/10.1101/gr.229202.

Kolmogorov, Mikhail, Jeffrey Yuan, Yu Lin, and Pavel A. Pevzner. 2019. “Assembly of Long, Error-Prone Reads Using Repeat Graphs.” Nature Biotechnology 37 (5): 540–46. https://doi.org/10.1038/s41587-019-0072-8.

Kozik, Alexander, Beth A. Rowan, Dean Lavelle, Lidija Berke, M. Eric Schranz, Richard W. Michelmore, and Alan C. Christensen. 2019. “The Alternative Reality of Plant Mitochondrial DNA: One Ring Does Not Rule Them All.” PLoS Genetics 15 (8): e1008373. https://doi.org/10.1371/journal.pgen.1008373.

Laslett, Dean, and Bjorn Canback. 2004. “ARAGORN, a Program to Detect tRNA Genes and tmRNA Genes in Nucleotide Sequences.” Nucleic Acids Research 32 (1): 11–16. https://doi.org/10.1093/nar/gkh152.

Li, Heng. 2013. “Aligning Sequence Reads, Clone Sequences and Assembly Contigs with BWA-MEM.” arXiv [q-bio.GN]. arXiv. http://arxiv.org/abs/1303.3997.

Li, Heng. 2016. “Minimap and Miniasm: Fast Mapping and de Novo Assembly for Noisy Long Sequences.” Bioinformatics 32 (14): 2103–10. https://doi.org/10.1093/bioinformatics/btw152.

Li, Heng. 2018. “Minimap2: Pairwise Alignment for Nucleotide Sequences.” Bioinformatics 34 (18): 3094–3100. https://doi.org/10.1093/bioinformatics/bty191.

Li, Heng, Bob Handsaker, Alec Wysoker, Tim Fennell, Jue Ruan, Nils Homer, Gabor Marth, Goncalo Abecasis, Richard Durbin, and 1000 Genome Project Data Processing Subgroup. 2009. “The Sequence Alignment/Map Format and SAMtools.” Bioinformatics 25 (16): 2078–79. https://doi.org/10.1093/bioinformatics/btp352.

McFadden, G. I. 1999. “Endosymbiosis and Evolution of the Plant Cell.” Current Opinion in Plant Biology 2 (6): 513–19. https://doi.org/10.1016/s1369-5266(99)00025-4.

McLay, Todd. n.d. “High Quality and High Yields of DNA for Long Read Sequencing Using a Sorbitol Prewash and CTAB or SDS.” Accessed April 21, 2020. https://www.genomicsforaustralianplants.com/wp-content/uploads/2020/03/DNA-extraction-Acacia-pycnantha.pdf.

O’Leary, Nuala A., Mathew W. Wright, J. Rodney Brister, Stacy Ciufo, Diana Haddad, Rich McVeigh, Bhanu Rajput, et al. 2016. “Reference Sequence (RefSeq) Database at NCBI: Current Status, Taxonomic Expansion, and Functional Annotation.” Nucleic Acids Research 44 (D1): D733–45. https://doi.org/10.1093/nar/gkv1189.

Rang, Franka J., Wigard P. Kloosterman, and Jeroen de Ridder. 2018. “From Squiggle to Basepair: Computational Approaches for Improving Nanopore Sequencing Read Accuracy.” Genome Biology 19 (1): 90. https://doi.org/10.1186/s13059-018-1462-9.

Sanchez-Puerta, M. Virginia, Alejandro Edera, Carolina L. Gandini, Anna V. Williams, Katharine A. Howell, Paul G. Nevill, and Ian Small. 2019. “Genome-Scale Transfer of Mitochondrial DNA from Legume Hosts to the Holoparasite Lophophytum Mirabile (Balanophoraceae).” Molecular Phylogenetics and Evolution 132 (March): 243–50. https://doi.org/10.1016/j.ympev.2018.12.006.

Sloan, Daniel B. 2013. “One Ring to Rule Them All? Genome Sequencing Provides New Insights into the ‘master Circle’model of Plant Mitochondrial DNA Structure.” The New Phytologist 200 (4): 978–85. https://nph.onlinelibrary.wiley.com/doi/abs/10.1111/nph.12395.

Tillich, Michael, Pascal Lehwark, Tommaso Pellizzer, Elena S. Ulbricht-Jones, Axel Fischer, Ralph Bock, and Stephan Greiner. 2017. “GeSeq - Versatile and Accurate Annotation of Organelle Genomes.” Nucleic Acids Research 45 (W1): W6–11. https://doi.org/10.1093/nar/gkx391.

Vaser, Robert, and Mile Šikić. 2020. “Raven: A de Novo Genome Assembler for Long Reads.” Cold Spring Harbor Laboratory. https://doi.org/10.1101/2020.08.07.242461.

Vaser, Robert, Ivan Sović, Niranjan Nagarajan, and Mile Šikić. 2017. “Fast and Accurate de Novo Genome Assembly from Long Uncorrected Reads.” Genome Research 27 (5): 737–46. https://doi.org/10.1101/gr.214270.116.

Walker, Bruce J., Thomas Abeel, Terrance Shea, Margaret Priest, Amr Abouelliel, Sharadha Sakthikumar, Christina A. Cuomo, et al. 2014. “Pilon: An Integrated Tool for Comprehensive Microbial Variant Detection and Genome Assembly Improvement.” PloS One 9 (11): e112963. https://doi.org/10.1371/journal.pone.0112963.

Wang, Weiwen, and Robert Lanfear. 2019. “Long-Reads Reveal That the Chloroplast Genome Exists in Two Distinct Versions in Most Plants.” Genome Biology and Evolution 11 (12): 3372–81. https://doi.org/10.1093/gbe/evz256.

Wang, Weiwen, Miriam Schalamun, Alejandro Morales-Suarez, David Kainer, Benjamin Schwessinger, and Robert Lanfear. 2018. “Assembly of Chloroplast Genomes with Long- and Short-Read Data: A Comparison of Approaches Using Eucalyptus Pauciflora as a Test Case.” BMC Genomics 19 (1): 977. https://doi.org/10.1186/s12864-018-5348-8.

Wheeler, Travis J., and Sean R. Eddy. 2013. “Nhmmer: DNA Homology Search with Profile HMMs.” Bioinformatics 29 (19): 2487–89. https://doi.org/10.1093/bioinformatics/btt403.

Wick, Ryan. n.d. “Filtlong.” Accessed April 21, 2020. https://github.com/rrwick/Filtlong.

Wick, Ryan R., and Kathryn E. Holt. 2019. “Benchmarking of Long-Read Assemblers for Prokaryote Whole Genome Sequencing.” F1000Research 8 (December): 2138. https://doi.org/10.12688/f1000research.21782.2.

Wick, Ryan R., Louise M. Judd, Claire L. Gorrie, and Kathryn E. Holt. 2017. “Unicycler: Resolving Bacterial Genome Assemblies from Short and Long Sequencing Reads.” PLoS Computational Biology 13 (6): e1005595. https://doi.org/10.1371/journal.pcbi.1005595.

Wick, Ryan R., Louise M. Judd, and Kathryn E. Holt. 2019. “Performance of Neural Network Basecalling Tools for Oxford Nanopore Sequencing.” Cold Spring Harbor Laboratory. https://doi.org/10.1101/543439.

Wick, Ryan R., Mark B. Schultz, Justin Zobel, and Kathryn E. Holt. 2015. “Bandage: Interactive Visualization of de Novo Genome Assemblies.” Bioinformatics 31 (20): 3350–52. https://doi.org/10.1093/bioinformatics/btv383.

Williams, Anna V., Laura M. Boykin, Katharine A. Howell, Paul G. Nevill, and Ian Small. 2015. “Correction: The Complete Sequence of the Acacia Ligulata Chloroplast Genome Reveals a Highly Divergent clpP1 Gene.” PloS One 10 (9): e0138367. https://doi.org/10.1371/journal.pone.0138367.

Williams, Anna V., Joseph T. Miller, Ian Small, Paul G. Nevill, and Laura M. Boykin. 2016. “Integration of Complete Chloroplast Genome Sequences with Small Amplicon Datasets Improves Phylogenetic Resolution in Acacia.” Molecular Phylogenetics and Evolution 96 (March): 1–8. https://doi.org/10.1016/j.ympev.2015.11.021.

Wu, Zhi-qiang, Xue-zhu Liao, Xiao-ni Zhang, Luke R. Tembrock, and Amanda Broz. 2020. “Genomic Architectural Variation of Plant mitochondria—A Review of Multichromosomal Structuring.” Journal of Systematics and Evolution 158 (August): 851. https://doi.org/10.1111/jse.12655.

Xu, Zhimeng, Yuting Mai, Denghui Liu, Wenjun He, Xinyuan Lin, Chi Xu, Lei Zhang, et al. 2020. “Fast-Bonito: A Faster Basecaller for Nanopore Sequencing.” Cold Spring Harbor Laboratory. https://doi.org/10.1101/2020.10.08.318535.

